# Exploiting allele-specific transcriptional effects of subclonal copy number alterations for genotype-phenotype mapping in cancer cell populations

**DOI:** 10.1101/2023.01.10.523464

**Authors:** Hongyu Shi, Marc J. Williams, Gryte Satas, Adam C. Weiner, Andrew McPherson, Sohrab P. Shah

## Abstract

Somatic copy number alterations drive aberrant gene expression in cancer cells. In tumors with high levels of chromosomal instability, subclonal copy number alterations (CNAs) are a prevalent feature which often result in heterogeneous cancer cell populations with distinct phenotypes^1^. However, the extent to which subclonal CNAs contribute to clone-specific phenotypes remains poorly understood, in part due to the lack of methods to quantify how CNAs influence gene expression at a subclone level. We developed TreeAlign, which computationally integrates independently sampled single-cell DNA and RNA sequencing data from the same cell population and explicitly models gene dosage effects from subclonal alterations. We show through quantitative benchmarking data and application to human cancer data with single cell DNA and RNA libraries that TreeAlign accurately encodes clone-specific transcriptional effects of subclonal CNAs, the impact of allelic imbalance on allele-specific transcription, and obviates the need to arbitrarily define genotypic clones from a phylogenetic tree *a priori*. Combined, these advances lead to highly granular definitions of clones with distinct copy-number driven expression programs with increased resolution and accuracy over competing methods. The resulting improvement in assignment of transcriptional phenotypes to genomic clones enables clone-clone gene expression comparisons and explicit inference of genes that are mechanistically altered through CNAs, and identification of expression programs that are genomically independent. Our approach sets the stage for dissecting the relative contribution of fixed genomic alterations and dynamic epigenetic processes on gene expression programs in cancer.

## INTRODUCTION

Genomic instability is a hallmark of human cancer which leads to copy number alterations (CNAs) in cancer cell genomes, and extensive intra-tumor heterogeneity^1–3^. It is well established that CNAs of driver oncogenes and tumor suppressors are causal determinants that change the fitness of cancer cells^4,5^, leading to clonal expansions, clone-clone variation^6^ and tumor evolution. Recent reports on the extent of cell-to-cell variation of CNAs in tumors (including in well understood oncogenes)^1^ raises the critical question of how granular subpopulations are phenotypically impacted by subclonal CNAs. Importantly, phenotypic impact of subclonal CNAs can have cell intrinsic effects and act as cell-extrinsic determinants of the tumor microenvironment^7^, further illustrating the importance of dissecting how CNAs modulate intra-tumor heterogeneity.

Previous studies using bulk sequencing techniques have investigated the association between clonal CNAs and gene expression^8–11^. The expression level of a gene can be influenced by copy-number dosage effects reflected by the significant positive correlation between gene expression and the underlying copy number (CN)^12^. However, gene dosage effects are not deterministic and may be subject to compensatory mechanisms, rendering the impact of CNAs on expression as highly variable across the genome. Transcriptional adaptive mechanisms^13^ including epigenetic modifications and downstream transcriptional regulation, can modulate copy number dosage effects^14–16^, further obscuring the direct impact of gene dosage. For example, the expression of certain immune response pathways often exhibit both CNA-dependent and CNA-independent expression^8^.

Theoretically, measuring single cell RNA and DNA data should elucidate how genotypes translate to phenotypes at single cell resolution. Technologies that sequence both RNA and DNA modalities from the same cell would be ideal for linking genomic alterations to transcriptional changes in tumor evolution. However, pioneering technologies^17,18^ have had limited throughput, lower quality and are still not mature enough for large-scale profiling of cancer cells. Sequencing single cell RNA or DNA independently allows more cells to be profiled and reveals a more comprehensive view of the cell populations, but requires computational integration of the two data modalities.

Several computational methods have been proposed for joint analysis of single cell DNA and RNA data. CloneAlign^19^ is a probabilistic framework to assign transcriptional profiles to genomic subclones based on the assumption that the expression level of a gene is proportional to its underlying copy number. More recent methods SCATrEx^20^ and CCNMF^21^ are also based on this assumption but use different methods to model the similarity between copy number profiles and gene expression patterns. However, these methods do not consider the possibility that transcriptional effects of copy number could be variable between genes and therefore lack the specificity to decipher genes that may be subject to dosage effects from those that are independent of CNAs. In addition, these methods require using predefined subclones from scDNA data as input which may propagate errors of uninformative subclones or may miss more granular gene dosage effects. More importantly, the revelation of phenotypic plasticity as a driver of genetically independent transcription in cancer cells^22–24^ motivates the need to disentangle genetic from epigenetic cell-to-cell variation. No available methods directly model dosage effects of subclonal CNAs, which is critical to infer which genes are deterministically modulated by subclonal CNAs and which genes are independent of CNAs. Moreover, recent advances have illuminated the extent to which allele-specific copy number alterations can mark clonal haplotypes both in DNA-based^1^ and RNA-based^25^ single cell analysis, illustrating both a methodological gap and analytical opportunity for integration.

In this study, we address the questions of how subclonal CNAs drive phenotypic divergence and evolution in cancer cells, and quantitatively encode (allele specific) copy number dosage effects in this process. We present a new method, TreeAlign, to enumerate and define CNA-driven clone-specific phenotypes, and also a statistical framework to compare the transcriptional readouts of genomically defined clones. TreeAlign is a Bayesian probabilistic model that maps gene expression profiles from scRNA to phylogenies from scDNA which i) obviates the need to identify clones *a priori* from a tree, ii) explicitly models dosage effects of each gene and iii) models allele-specific CNAs to better resolve clonal mappings.

Through extensive simulation, we demonstrate that the TreeAlign outperforms alternative approaches in terms of clone assignment and gene dosage effect prediction. Applying TreeAlign to both primary tumors and cancer cell lines, we characterized the phenotypic differences between tumor subclones, investigated the contribution of subclonal CNAs to clone-specific gene expression patterns in cancer cells and identify common expression programs which are altered by subconal CNAs.

## RESULTS

### TreeAlign: a probabilistic graphical model for clone assignment and dosage effect inference

We developed TreeAlign, a probabilistic graphical model of scRNA transcriptional profiles mapped to a scDNA-derived phylogenetic tree **(Fig.1)**. The model jointly infers clone assignments, clone-specific copy number dosage effects and optionally, models clone-specific allelic transcriptional effects. The TreeAlign framework assumes that there exists a subset of genes whose expression is positively correlated with the underlying copy number. For each gene, the correlation between subclonal CNAs and gene expression is modeled by *k*, where *k*∈ {0,1}**(Fig. 1c)** is a switching indicator variable such that the probability *p*(*k* =1) represents the probability of a gene with clone-specific copy number dosage effects. As such, genes without dosage effects will have low *p*(*k*) and will not contribute to the clone assigning process. To infer clone assignments and *p*(*k*), TreeAlign requires three inputs: 1, a cell × gene matrix of raw read counts from scRNA-seq, 2. a cell × gene copy number matrix estimated from scDNA-data and 3. A phylogenetic tree (or optionally, predetermined clone labels) for scDNA profiles. TreeAlign can either assign expression profiles to predefined clone labels, similar to CloneAlign^19^ or operate on a phylogenetic tree directly and assign cells to clades of the phylogeny **(Fig. 1a)**. When TreeAlign takes a phylogenetic tree as input, it applies a Bayesian hierarchical model recursively starting from the root of the phylogenetic tree and computes the probability that expression profiles in scRNA can be mapped to a subtree. When the genomic or phenotypic differences between two subtrees become too small to allow confident assignment of expression profiles, TreeAlign will stop its recursion and return the resulting subtrees.

**Fig. 1:**
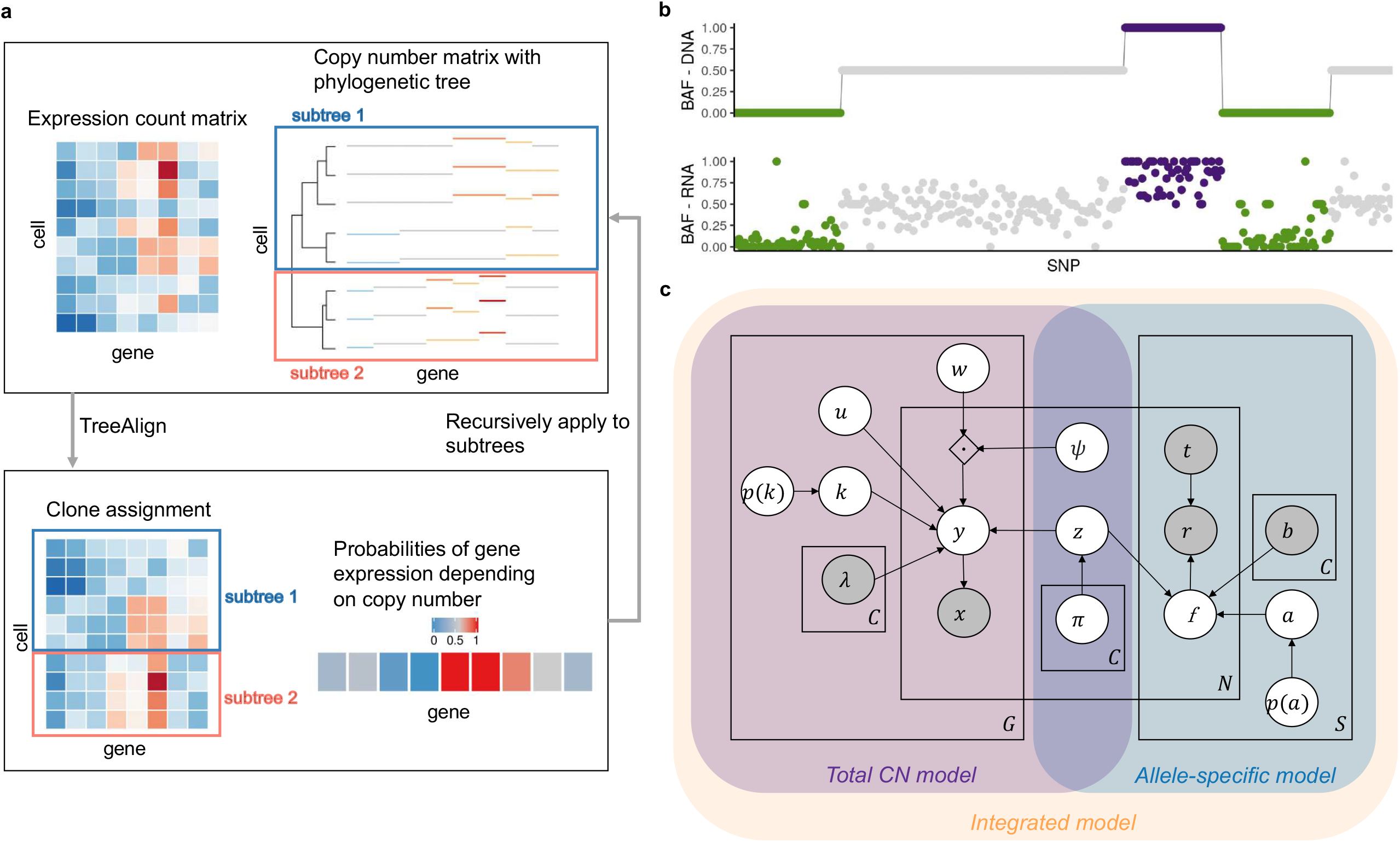
Overview of TreeAlign. **a**, TreeAlign takes raw count data from scRNA-seq, the copy number matrix and the phylogenetic tree from scDNA-seq. By recursively assigning the expression profiles to phylogenetic subtrees, TreeAlign infers the clone-of-origin of cells identified in scRNA-seq and the dosage effects of clone-specific copy number alterations. **b**, Allelic imbalance as measured by B allele frequency can be inferred from DNA-data and RNA-data. We assume a positive correlation between the two measurements to improve clone assignment. **c**, Graphical model of TreeAlign.

In addition to aberrant gene expression levels, allele-specific CNAs also lead to allele-specific expression imbalance which is detectable in scRNA data^26,27^ **(Fig. 1b)**. In particular, genomic segments harboring loss of heterozygosity deterministically leads to mono-allelic expression of genes in the segment. To exploit how allelic imbalance modulates allele specific expression, we extended TreeAlign to model both total CN and allelic imbalance data **(Fig. 1c, Extended Data Fig. 1)**. Given the B allele frequencies (BAFs) estimated from scDNA data haplotype blocks using SIGNALS^1^ and allele-specific expression at corresponding heterozygous SNPs in scRNA data, the allele-specific model contributes to clone assignment and infers the probability of the allele assignment *p*(*a* =1), *a* ∈ {0,1}which indicates whether the SNP is on allele B or not.

The software for TreeAlign (https://github.com/AlexHelloWorld/TreeAlign) is implemented in Python using Pyro and is publicly available. Our implementation allows users to run the total CN model, allele-specific model and integrated model by providing different inputs. See **Methods** for additional mathematical, inference and implementation details.

### Performance of TreeAlign on simulated data

We first evaluated TreeAlign on synthetic datasets, quantifying the effect of three main parameters in the input data: number of cells (100 - 5000), number of genes (100 - 1000) and proportions of genes with dosage effects (10%-100%). Simulations were performed using the generative model of CloneAlign^19^. We compared the performance of assigning expression profiles to ground truth predefined clones between TreeAlign, CloneAlign and InferCNV^28^. InferCNV was originally developed for inferring CNAs from gene expression data, but has also been repurposed for clone assignment in some studies^29^. InferCNV analysis in this context acts as a way of inferring clone assignment without the benefit of the scDNA data. Compared to CloneAlign and InferCNV, TreeAlign performed significantly better in terms of clone assignment accuracy especially in the regime where fewer genes exhibit dosage effects **(Fig. 2a, Extended Data Table 1)**. For example, in the regime of 60% of genes with dosage effects (1000 cells, 500 genes), TreeAlign achieved clone assignment accuracy of 91.1%, compared to CloneAlign with 75.1% accuracy. The improvement in clone assignment accuracy was consistent across all cell number and gene dosage effect simulation scenarios **(Extended Data Fig. 2a)**. We also tested performance with phylogenetic tree inputs to evaluate if TreeAlign could achieve similar results on tree input compared to pre-defined clone input. Similar to the ‘clone’ regime, these simulations varied the proportion of genes with gene dosage effects in 10% increments. TreeAlign was able to assign expression profiles back to the corresponding clades of the phylogeny with similar accuracies compared to the clone input in regimes with >40% genes with dosage effects **(Fig. 2b, Extended Data Fig. 2b)**. Together these evaluations reflect that the model effectively obviates *a priori* tree cutting without paying a penalty in accurate clone mapping.

**Fig. 2:**
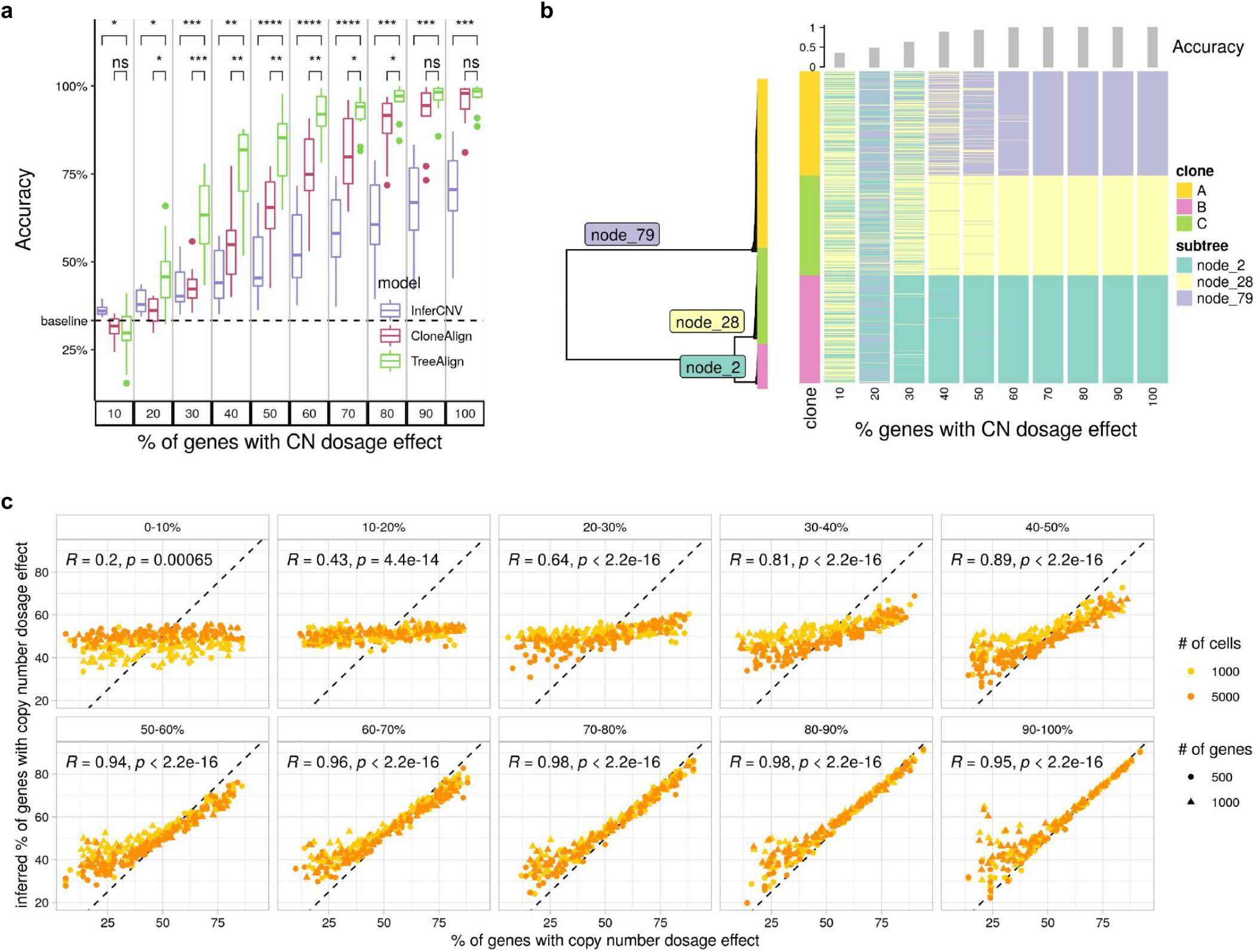
Performance of TreeAlign on simulated data. **a**, Clone assignment accuracy of TreeAlign, CloneAlign and InferCNV on simulated datasets (500 cells, 1000 genes, 3 clones) containing varying proportions of genes with copy number dosage effects. *P<0.05, **P<0.01, ***P<0.001, ****P<0.0001. Brackets: Wilcoxon signed-rank test. **b**, Phylogenetic tree (left) of cells from patient 081 constructed using scDNA-data. Heat map (right) of clone assignment by TreeAlign. Each column shows the assignment of simulated expression profiles to subtrees of the phylogeny. The bar chart above shows the overall accuracy of clone assignment. **c**. Scatter plots comparing inferred gene dosage effect frequencies and the simulated frequencies. Each panel groups genes with similar expression levels from low expression genes (0-10%) to high expression genes (90-100%). Pearson correlation coefficients (R) and P values for the linear fit are shown.

We also evaluated the accuracy of predicting dosage effects for each gene in the input datasets. We compared the simulated and predicted (using *p*(*k*) as an estimate) frequency of genes with CN dosage effects. For high expression genes, simulated and predicted frequencies were highly concordant **(Fig. 2c)**. For datasets with >=50% of genes with dosage effects, the mean area under the receiver-operator curve (AUC) was >=0.99 for genes with relatively high expression level (genes in top 40% in terms of normalized expression levels) **(Extended Data Fig. 3)**. This establishes *p*(*k*) as an accurate representation of gene dosage effects and the ability to distinguish genes with dosage effects from those without dosage effects.

### TreeAlign assigns HGSC expression profiles to phylogeny accurately

We next investigated TreeAlign’s performance on real-world patient derived data from high grade serous ovarian cancer (HGSC). We first applied TreeAlign on single cell sequencing data from a HGSC patient (patient 022)^7^. Tumor samples were obtained from both left and right adnexa sites of the patient. scDNA (n = 1050 cells) and scRNA (n = 4134 cells) data were generated through Direct Library Preparation (DLP+)^30^ and 10X genomics single-cell RNA-seq^31^ respectively. 3579 (86.6%) ovarian cancer cells profiled by scRNA were assigned to 4 subclones identified by scDNA-seq. The expression profiles of clone C and D are overlapped on the UMAP embedding, while separated from the profiles of clone A and clone B, which coincides with the shorter phylogenetic distance between clone C and D **(Fig. 3a)**. The separation of cells by assigned clones on the expression-based UMAP also suggests that the genetic subclones possess distinct transcriptional phenotypes.

**Fig. 3:**
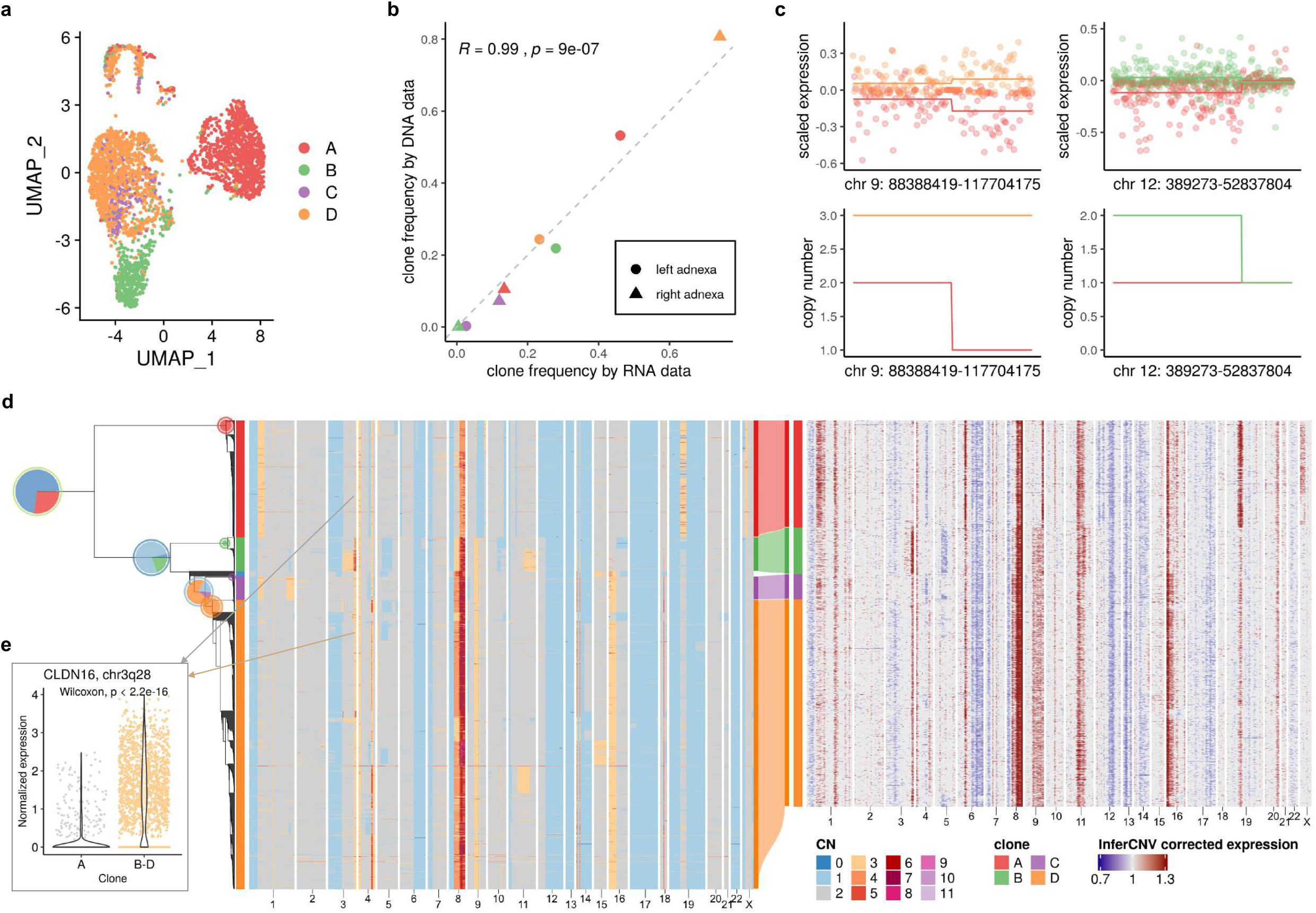
TreeAlign assigns HGSC expression profiles to phylogeny accurately. **a**, UMAP plot of scRNA-data from patient 022 colored by clone labels assigned by TreeAlign. **b**, Correlation between clone frequencies of patient 022 estimated by scRNA-data (x axis) and scDNA-data (y axis). **c**, Scaled expression and copy number profiles for regions on chromosome 9 and 12 as a function of genes ordered by genomic location. **d**, Single cell phylogenetic tree of patient 022 constructed with scDNA-data (left). Pie charts on the tree showing how TreeAlign assigns cell expression profiles to subtrees recursively. The pie charts are colored by the proportions of cell expression profiles assigned to downstream subtrees. The outer ring color of the pie charts denotes the current subtree. Left heat map, total copy number from scDNA; right heat map, InferCNV corrected expression from scRNA; middle Sankey chart, clone assignments from RNA to DNA. **e**, Normalized expression of CLDN16 in clone A and clone B - D.

We confirmed the clone assignment accuracy of TreeAlign by comparing the clonal frequencies estimated by RNA and DNA data **(Fig. 3b)**. As both scRNA and scDNA data were generated by sampling from the same populations of cells, the clonal frequency estimated by the two methods should be consistent. Clonal frequencies in the left and right adnexa sample from the two modalities were significantly correlated (R = 0.99, P = 9 × 10^−7^). In addition, copy number alterations inferred for scRNA cells using InferCNV^28^ were concordant with the scDNA based CNA of the clones to which scRNA cells were assigned **(Fig. 3d)**. For example, notable clone specific copy number changes can be seen in both scDNA and scRNA on chromosome X in clone A. Clone B specific amplification on 3q, Clone C and Clone D specific amplification on 16p can also be observed in both scDNA and scRNA. By comparing the RNA-derived copy number profiles with scDNA data, we noticed that inferring copy number from RNA data is not always accurate. For example, the inferred profiles missed the focal amplification on chromosome 18. We also held out genes from chromosome 9 and chromosome 12 and re-ran TreeAlign with the remaining genes. 98.8% cells were assigned consistently as compared to results using the full dataset. Clone level gene expression on chromosome 9 and 12 was consistent with the corresponding copy numbers **(Fig. 3c)**. These results demonstrated a proof of principle that TreeAlign can properly integrate scRNA and scDNA datasets and highlighted that scDNA-seq can provide valuable information on CNAs and tumor subclonal structures which would be difficult to detect with expression data only.

We also applied TreeAlign to previously published data from a gastric cell line NCI-N87 generated by 10x genomics single-cell CNV and 10x scRNA assays^32^. TreeAlign assigned 3212 cells from scRNA to three clones identified in scDNA. The clonal frequencies estimated by both assays were closely aligned **(Extended Data Fig. 4)**. As for the patient 022 data, the scRNA cells showed subclonal copy number similar to the scDNA clones to which they were assigned, thus illustrating that TreeAlign also performs well with 10x scDNA data.

### Incorporating allele specific expression increases clone assignment resolution

We next investigated the extent to which accurate clone assignment solely based on allele specific expression could be performed. We inferred allele specific copy number and BAF using scDNA data from patient 022 with SIGNALS^1^. The allele specific heat map **(Fig. 4a)** revealed characteristic patterns of clonal loss of heterozygosity in whole chromosomes (e.g. chr 6,13, 14, 17) but also subclonal losses (e.g. chr 9q in clone A and parallel losses on chr 5 across multiple subclones). With the allele-specific model, we assigned cells from scRNA to clone A as identified by scDNA in patient 022. Clone assignments were consistent between the allele specific model and the total CN model with 87% cells concordant. The clone-specific BAF estimated from scRNA accurately reflected scDNA **(Extended Data Fig. 6a)**, with the exception of SNPs on chromosome X which showed allelic imbalance in scRNA but not in scDNA due to X-inactivation. The predicted allele assignments of SNPs from the allele-specific model were also consistent with haplotype phasing from scDNA (AUC=0.84) **(Fig. 4f)**. These results suggest that allelic imbalance information can be effectively exploited for clonal mapping.

**Fig. 4:**
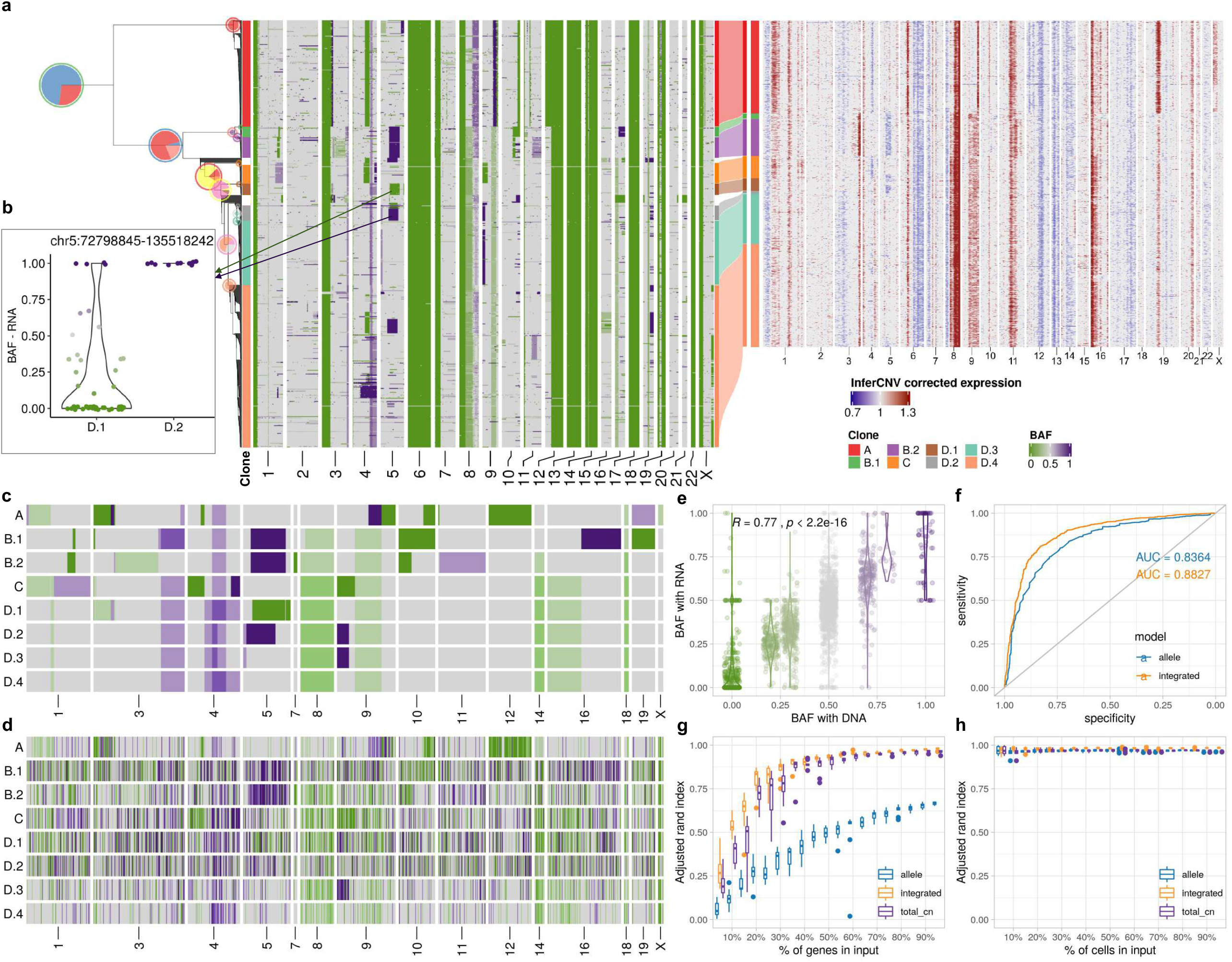
Incorporating allele specific expression increases clone assignment resolution. **a**, Integrated TreeAlign model assigns expression profiles to phylogeny of patient 022. Left heat map, single cell BAF profiles estimated from scDNA-data using SIGNALS, annotated with clone labels on the left side (BAF profiles without clone label represent cells ignored by TreeAlign) (Methods). **b**, BAF estimated from scRNA in clone D.1 and D.2 at region chr5:72,798,845-135,518,242. **c-d**, BAF of subclones with (c) scDNA and (d) scRNA. **e**, Correlation between BAF estimated with scRNA and BAF estimated with scDNA in patient 022. Annotations at the top indicate the Pearson correlation coefficient (R) and P value derived from a linear regression. **f**, ROC curves for predicting *p*(*a* =1) with allele-specific TreeAlign and integrated TreeAlign. **g**, Robustness of clone assignment to gene subsampling in patient 022. Adjusted rand index was calculated by comparing clone assignments using subsampled datasets to the complete dataset. **h**, Robustness of clone assignment to cell subsampling in patient 022.

We then applied the integrated model utilizing both total CN and allele-specific information on data from patient 022. Relative to the total CN model, the integrated model mapped scRNA cells to smaller subclones **(Fig. 4a)**. Specifically we note when considering allele specificity, Clone B was subdivided into two subclones (B.1 and B.2). Clone B.1 had an additional deletion at 16q leading to LOH and a gain of 10q leading to allelic imbalance, whereas Clone B.2 had an amplification at 11q with increased BAF **(Fig. 4a)**. Clone D was further divided into four subclones (D.1, D.2, D.3 and D.4). Clone D.1 and clone D.2 both had a deletion on chromosome 5, but the deletion events occurred on different alleles in the two subclones with different breakpoints, each of which was distinct from the 5q deletion on Clone B, indicative that parallel evolution is indeed reflected in transcription with the allele specific model **(Fig. 4b)**. We also estimated BAF for each of the subclones assigned from the scRNA data. Subclonal BAF estimated with scRNA and scDNA data were significantly correlated (0.25 < R < 0.53 for each subclone, P < 2.2 × 10^−22^) **(Fig. 4e; Extended Data Fig. 6c)**, consistent with more accurate clone assignment. With integrated TreeAlign, we also achieved better performance for predicting allele assignments of SNPs compared to the allele-specific model **(Fig. 4f)**. We note that recent identifications of parallel allelic-specific alterations whereby maternal and paternal alleles are independently lost or gained in different cells^26,27,33^ would further complicate clonal mapping, if allele specificity is not taken into account. Here we show that mono-alleleic expression of maternal and paternal alleles is consistent with coincident maternal and paternal allelic loss in different clones **(Fig. 4b)**. The allele-specific TreeAlign model correctly assigns cells at this level of granularity that would otherwise be missed.

We compared the performance of total CN, allele-specific and integrated TreeAlign using subsampled datasets of patient 022 and evaluating against results from the full dataset. All three models were robust to reduced numbers of cells **(Fig. 4h, Extended Data Table 2)**. The integrated model performed significantly better when fewer genomic regions were included in the input suggesting it is more robust when there are few copy number differences between subclones **(Fig. 4g)**, and the allele-specific model without total CN is inferior, as expected.

### Inferring copy number dosage effects in human cancer data

We next compared the integrated model to the total CN model on a recently published cohort of cell lines and primary tumors with scDNA and scRNA matched data from Funnell et al.^1^ We applied TreeAlign on data previously collected from patient derived xenografts of TNBC (n = 2), HGSC (n = 7), and from primary ovarian cancer (n = 1). In addition we tested the model on 184-hTERT (n = 6) cell lines engineered to induce genomic instability from a diploid background with CRISPR loss of function of *TP53* combined with *BRCA1* or *BRCA2*. Both integrated and total CN TreeAlign were run on matched DLP+ and 10x scRNA-seq data. In this analysis, we investigated the impact of *p*(*k*) on interpretability of genotype-phenotype linking. As expected, the integrated model characterized more clones **(Fig. 5b)** and achieved a lower number of cells not confidently assigned to a subclone **(Fig. 5c)**. For cells that were assigned confidently by the integrated model but not the total CN model, their InferCNV corrected expression showed higher correlation coefficient with the CN profiles of subclones assigned by the integrated model compared to random subclones **(Fig. 5d; Extended Data Fig. 7)**, implying better performance of the integrated model.

**Fig. 5:**
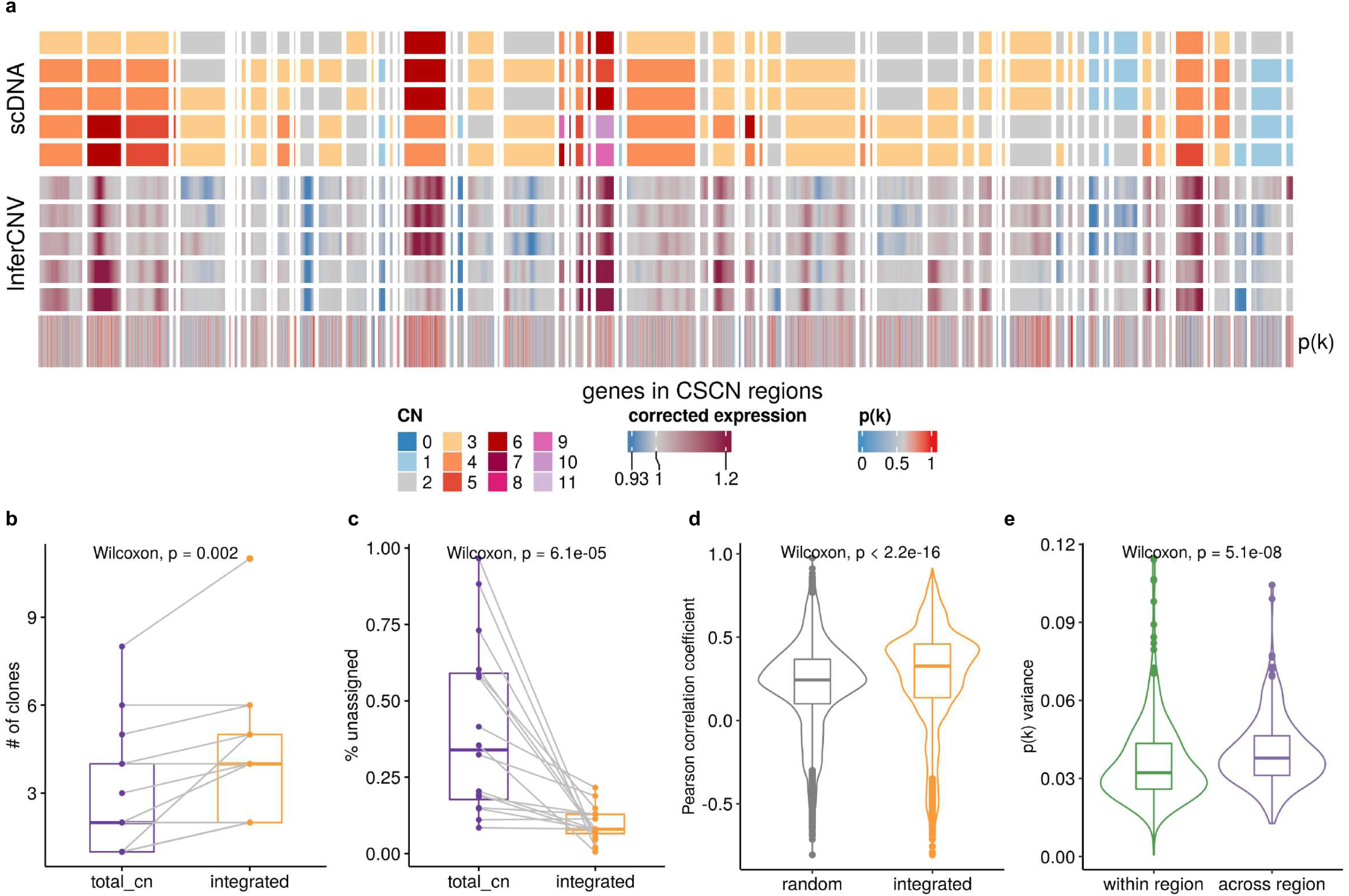
Inferring copy number dosage effects in human cancer data. **a**, Heat map representations of genes in CSCN regions in HGSC sample SA1096. Top heat map: clone-level total copy number from scDNA; bottom heat map: InferCNV corrected expression profiles from scRNA; bottom track: p(k) estimated by TreeAlign. **b**, Number of clones characterized by total CN and integrated model (Wilcoxon signed-rank test). **c**, Frequencies of unassigned cells (Methods) from total CN and integrated model (Wilcoxon signed-rank test). **d**, Distribution of Pearson correlation coefficients (R) between scDNA estimated total copy number and InferCNV corrected expression for unassigned cells from total CN model. Left, correlation distribution calculated by comparing InferCNV profiles to CN profiles of a random subclone; Right, correlation distribution calculated by comparing InferCNV profiles to CN profiles of subclones assigned by integrated TreeAlign. **c**, Variance of p(k) sampled from the same genomic regions and across regions.

For high expression genes (top 40% in terms of normalized expression levels) located in clone specific copy number (CSCN) regions, 77.3% had *p*(*k*) > 0.5 suggesting their expression is dependent on copy number **(Extended Data Fig. 8a, b, c)**. When we summarized *p*(*k*) by genomic locations, we noticed that genes located at the same CSCN region had more consistent *p*(*k*). Notably, *p*(*k*) of genes in a contiguous region exhibited significantly lower variation compared to randomly sampled genes across different regions **(Fig. 5a, e)**. This is consistent with multiple genes in a CNA transcriptionally impacted by a singular genomic event. In addition to broad regions of the genome, we note that subclonal high-level amplifications affecting known oncogenes have been identified previously^1^. Using TreeAlign, we also identified subclonal amplifications of oncogenes accompanied by consistent changes in gene expression. For example, in SA1096 and OV2295, subclonal upregulation of MYC expression coincides with the clone-specific MYC amplification with *p*(*k*) > 0.8 **(Extended Data Fig. 9a)**. To investigate whether MYC pathway activation was also impacted by non-CNA driven effects, we performed pathway enrichment on genes with low *p*(*k*) and found genes in the Hallmark MYC Target V1 gene set^34^ in OV2295, SA1052 and SA610. Combined with HLAMP results, this suggests the pathway can be regulated by both CN dosage effects and other (potentially non-genomic) effects at the subclonal level **(Extended Data Fig. 9b, c)**, further highlighting the importance of p(k) for interpreting the mechanism of gene dysregulation.

### Clone-specific transcriptional profiles highlight clonal divergence in immune pathways

We next sought to interpret clone-specific transcriptional phenotypes and phenotypic divergence during clonal evolution from TreeAlign mappings. For patient 022, differential expression and gene set enrichment analysis identified genes and pathways upregulated in each clone **(Fig. 6a, b)**. In total, we found 1346 genes significantly upregulated (adjusted P < 0.05, MAST^35^) in at least one of the subclones in patient 022. 52.1% (701) of these genes were not located in CSCN regions, while 47.9% (645) genes were located within CSCN regions. For 90.7% (585/645) of genes in CSCN regions, *p*(*k*) was > 0.5, reflecting probable gene dosage effects.

**Fig. 6:**
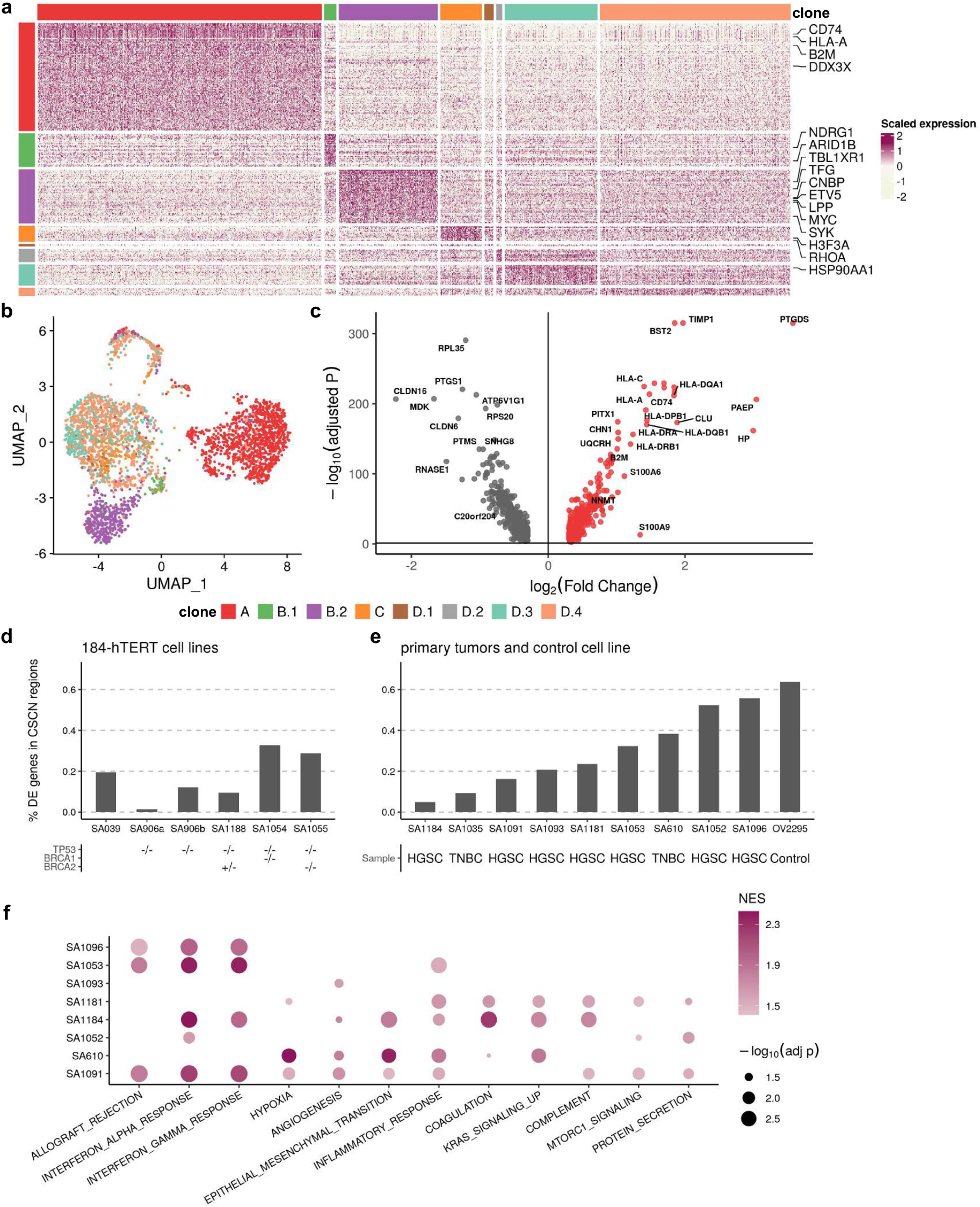
Clone-specific transcriptional profiles highlight clonal divergence in immune pathways. **a**, Scaled expression of upregulated genes in each subclone in patient 022, showing genes in rows and subclones in columns. Genes in the COSMIC Cancer Gene Census42 are highlighted. **b-c**, Proportions of **f** subclonal differentially expressed genes located in CSCN regions for (b) 184-hTERT cell lines, (c) an HGSC control cell line and primary tumors. **d**, UMAP embedding of expression profiles from patient 022 colored by clone labels assigned by integrated TreeAlign model. **e**, Differentially expressed genes between clone A and other subclones (clone B - D) in patient 022. **f**, Pathways with clone-specific expression patterns in TNBC and HGSC tumors.

Immune related pathways such as IFN-α and IFN-γ response were differentially expressed, and with increased relative expression in clone A **(Fig. 6c, Extended Data Fig. 11e and Extended Data Table 3)**. Clone A contains cells from both right and left adnexa, thus dysregulation of these pathways cannot be simply explained by the microenvironment of clone A. Differential expression of immune related pathways was also found between more closely related subclones. Compared to clone B.2, clone B.1 also has enriched expression in IFN-α and IFN-γ signaling pathways and downregulation in MYC targets V1 and G2M checkpoint gene sets **(Extended Data Fig. 10a; Extended Data Fig. 11b)**. Clone D.4, compared to other clone D subclones, had down-regulated TNF-α signaling via NFκB **(Extended Data Fig. 10b, f; Extended Data Fig. 11c)**. Seeking to explain the relative contribution of subclonal CNAs to differentially expressed pathways, we analyzed the proportion of differentially expressed genes found in subclonal CNAs for each pathway. Only 17.4% (4/23) of differentially expressed genes in the Allograft Rejection gene set are in CSCN regions compared to 61.5% (24/39) in the MYC Targets V1 gene set highlighting the distinct impact of subclonal CNA between pathways **(Extended Data Fig. 10h)**.

We conducted a similar analysis on data from Funnell et al. Differential expression analysis revealed varying proportions of DE genes located in CSCN regions ranging from 1.3% to 63.9%, indicating that transcriptional heterogeneity due to cis-acting subclonal CNAs varied across tumors **(Fig. 6d, e)**. In addition to pathways such as *KRAS* signaling and EMT which are known to be important in these tumors, IFN-α and IFN-γ response pathways also show frequent variable expression between subclones of primary TNBC and HGSC **(Fig. 6f)**. IFN signaling has important immune modulatory effects, and has been previously linked to immune evasion and resistance to immunotherapy^36^. The recurrent differential expression of immune related pathways between subclones suggests their importance in clonal divergence in these cancers of genomic instability.

## DISCUSSION

TreeAlign establishes a probabilistic framework for integration of scRNA and scDNA data and inference of dosage effects of subclonal CNAs. TreeAlign achieves high accuracy of assigning single cell expression profiles to genetic subclones and was built to operate on phylogenetic trees directly, therefore informing phenotypically divergent subclones during the recursive clone assignment process. In addition to scRNA and scDNA integration, TreeAlign also disentangles the *in cis* dosage effects of subclonal CNAs which highlights highly regulated pathways in clonal evolution. The model has improved flexibility allowing either total or allelic copy number or both to be used as input. With additional allele-specific information, TreeAlign has improved prediction accuracy and model robustness and is able to identify more refined clonal structure.

We expect potential extensions of TreeAlign for integration of other single cell data modalities such as single-cell epigenetic data. Current methods for integration of scRNA and scATAC data are primarily based on nearest neighbor graphs or other distance metrics to match similar cells across multimodal datasets^37^. The advantage of TreeAlign is that it estimates how well the expression of a gene matches with the given biological assumption, hence it is more interpretable and provides explanations for gene expression variations.

The emergence of more single cell multimodal datasets enable future studies to further reveal how genotypes translate to phenotypes and how ongoing mutational processes drive clonal diversification and evolution in cancer cells. It remains an open question whether the CN-expression relation is consistent across tumors and whether application at scale can reveal phenotypic consequences of copy number alterations at subclonal resolution. Furthermore, as TreeAlign also integrates allele-specific CN and expression, it would be interesting to investigate patterns of LOH and allele-specific expression on a subclone level as modulators of germline alterations and bi-allelic inactivation to better understand these events in the context of tumor heterogeneity and clonal evolution. We expect that concepts introduced in TreeAlign will facilitate the integration of single cell multimodal datasets and the interpretation of associations between modalities.

In conclusion, we anticipate that studying how copy number alterations impact gene expression programs in cancer applies broadly to different questions in cancer biology including etiology, tumor evolution, drug resistance and metastasis. In these settings, TreeAlign provides a flexible and scalable method for explaining gene expression with subclonal CNAs as a quantitative framework to arrive at mechanistic hypotheses from multimodal single cell data. Our approach provides a new tool to disentangle the relative contribution of fixed genomic alterations and other dynamic processes on gene expression programs in cancer.

## METHODS

### The TreeAlign Model

#### Model description and inference

The TreeAlign model is a probabilistic graphical model as shown in Fig. 1c. Here we describe the model in detail and how the model is fit to data. Let *X* be a cell×gene expression matrix of raw counts from scRNA-seq for *N* cells and *G* genes. Let *λ* be a gene×clone copy number matrix for *G* genes and *C* clones. To assign cells from the expression matrix to a clone in copy number matrix, we use a categorical vector *z* = {*z*_*n*_} which indicates the clone to which a cell should be assigned

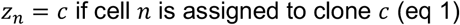

To indicate whether the expression of a gene *G* is dependent on underlying copy number, we introduce another indicator vector *k* = {*k*_*g*_}

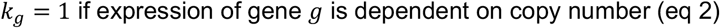

Our assumption is that *y*_*ng*_ - the expected expression of gene *g* in cell *n* - will be proportional to the copy number of gene *g* in clone *c* to which cell *n* is assigned, if expression of gene *g* is dependent on copy number as indicated by *k*_*g*_. Based on this assumption, our model is:

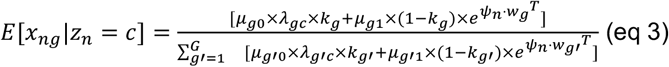

where *μ*_*g*0_ is the per-copy expression of gene *g* if the expression is dependent on copy number while *μ*_*g*1_ is the expression of gene *g* if its expression is independent of copy number. The intuition is when *k*_*g*_ = 1, we expect the expression of *g* is proportional to its copy number; when *k*_*g*_ = 0, the expression of *g* is not dependent on the underlying copy number. The inner product *ψ*_*n*_ · *w*_*g*_^*T*^ introduces noise into the model to avoid overfitting. We specified a softplus-Normal prior over the per-copy expression *μ*_*g*0_ and *μ*_*g*1_. Multinomial likelihood was used to model the raw count from scRNA with a mean given by (eq 3). Detailed definitions and distribution assumptions of random variables and data are described in Extended Data Fig. 1.

Inference is performed using stochastic variational inference in the Pyro package. We generate the variational distributions using the AutoDelta function which uses Delta distributions to construct a MAP guide over the latent space. Optimization is performed using the Adam optimizer. By default, we set a learning rate of 0.1 and the convergence is determined when the relative change in ELBO is lower than 10^−5^ by default.

#### Incorporating phylogeny as input

In addition to the gene×clone copy number matrix, TreeAlign can also take the cell×gene copy number matrix from scDNA directly along with the phylogenetic tree constructed from this matrix as input. Starting from the root of the phylogeny, TreeAlign summarizes the copy number of gene *g* for each clade by taking the mode of copy number, and assigns cells from scRNA to clade-level CN profiles. This process is repeated recursively from the root of the phylogeny to smaller clades until: i) TreeAlign can no longer assign cells consistently in multiple runs (less than 70% cells have consistent assignments between runs by default), or ii) the number of genes located in CSCN regions becomes too small (100 genes in CSCN regions by default), or iii) Limited number of cells remain in scDNA or scRNA (100 by default). By default, TreeAlign also ignores subclades with less than 20 cells in scDNA. Some scRNA cells may remain unassigned to the scDNA phylogenetic tree. For a single cell, if the clone assignment probability *π*_*c*_ < 0.8 or clone assignments are not consistent in 70% of repeated runs, the cell will be denoted as unassigned. This feature is important to the model because there might be incomplete sampling of a given tumor, leading to a subclone only appearing in one of the two data modalities. Note, all parameters are fully configurable at run time by the user.

#### Incorporating allele-specific information

To use allele specific copy number information for clone assignment, we set up a separate model - allele-specific TreeAlign which only takes in allele specific information. The input to allele-specific TreeAlign includes single cell level B allele frequencies at heterozygous SNPs estimated from scDNA-data and read counts of reference allele and alternative allele of these SNPs from scRNA-data. The underlying assumption is that the allelic imbalance in the genome is positively correlated to the imbalanced expression from reference allele and alternative allele as observed with scRNA-seq. To indicate whether the B allele defined with scDNA-data is the reference allele in gene expression data, we introduce an optional indicator variable *a*_*g*_.

*a*_*g*_ = 1 if B allele defined in scDNA is the reference allele in scRNA

The integrated TreeAlign model was constructed by combining the total-CN model and the allele-specific model.

#### Benchmarking clone assignment and dosage effect prediction with simulations

Simulations were performed similarly as described previously^19^. CloneAlign v.2.0 model was fit to the MSK-SPECTRUM patient 081 dataset to obtain the empirical estimations of model parameters. Then we simulated from CloneAlign considering the following scenarios: 1. Varying proportion (10%, 20%, 30%, …, 90%) of genes with dosage effect. 2. Varying number of genes (100, 500 and 1000) in CSCN regions. 3. Varying number of cells (100, 1000 and 5000) in scRNA.

We compared TreeAlign to CloneAlign and InferCNV v.1.3.5 in terms of the performance of clone assignment. For CloneAlign, we summarized clone-level copy number by calculating the mode of copy number for each gene and ran CloneAlign with default parameters. For InferCNV, we used the recommended setting for 10X. 3,200 non-cancer cells were randomly sampled from the SPECTRUM dataset and used as the set of reference “normal” cells. To assign clones with InferCNV, we calculated Pearson correlation coefficient between InferCNV corrected gene expression profile (expr.infercnv.dat) and the clone-level copy number profiles from scDNA. Cells from scRNA-seq were assigned to the clone according to the highest correlation coefficient. Accuracy of clone assignment was computed to compare the performance of the three methods. We also evaluated the TreeAlign’s performance on predicting CN dosage effects. We calculated the area under the curve (AUC) using *p*(*k*) output by TreeAlign.

#### MSK SPECTRUM data

We obtained matched scRNA and scDNA from two HGSC patients (patient 022 and patient 081) from the MSK SPECTRUM cohort^7^. Samples were collected under Memorial Sloan Kettering Cancer Center’s institutional IRB protocol 15-200 and 06-107. Single cell suspensions from surgically excised tissues were generated and flow sorted on CD45 to separate the immune component as previously described ^7^. CD45 negative fractions were then sequenced using the DLP+ platform as previously described ^1,30,38^.

#### Gastric cancer cell line data

Preprocessed scDNA data and scRNA count matrix of the gastric cancer cell line (NCI-N87)^32^ were downloaded from SRA (PRJNA498809) and GEO (GSE142750). Copy number calling for scDNA were performed using the Cellranger-DNA pipeline using default parameters.

#### HGSC, TNBC and additional cell line data

scRNA and scDNA from 7 primary HGSC (SA1093, SA1052, SA1053, SA1181, SA1184, SA1091, SA1096), 2 primary TNBC (SA1035, SA610), 1 ovarian cancer cell line (OV2295) and 6 hTERT-184 cell lines (SA039, SA1054, SA1055, SA1188, SA906a, SA906b) were obtained and processed as described previously^1^.

#### scDNA data analysis

scDNA DLP+ data was processed as previously described^1,30^. Cells with quality score > 0.75 and not in S-phase were retained for downstream analysis. Allele specific copy number was called using SIGNALS^1^, which provides allele specific copy number of the from A|B in 500kb bins across the genome. A and B being the copy number of alleles A and B respectively with *total CN* = *A* + *B*. As the single cell data is sparse, only a subset of germline SNPs have coverage in each cell, therefore to produce the input required for TreeAlign (B-Allele frequencies per SNP per cell), we impute the BAF of each SNP assuming that a SNP will have the same BAF as the bin in which the SNP resides.

#### Clustering and phylogenetic inference

Clustering and phylogenetic inference of scDNA was performed using UMAP and HDBSCAN (parameters min_samples = 20, min_cluster_size = 30, cluster_selection_epsilon = 0.2). For patient 022, we also constructed phylogenetic trees using Sitka^38^ as previously described.

#### Genotyping SNPs in scRNAseq cells

SNPs identified in scDNA-seq and matched bulk whole genome sequencing were genotyped in each single cell using cell-snplite^39^ with default parameters.

#### scRNA data analysis

scRNA data were processed as previously described^7^. Read alignment and barcode filtering were performed by CellRanger v.3.1.0. Cancer cell identification was performed with CellAssign. Principal-component analysis (PCA) was performed on the top 2000 highly variable features output by function FindVariableFeatures using Seurat v.4.2^40^. UMAP embeddings and visualization were generated using the first 20 principal components. Unsupervised clustering was performed using FindNeighbors function followed by FindClusters function (resolution = 0.2).

#### Differential expression and gene set enrichment analysis

Differential expression analysis was performed using FindAllMarkers and FindMarkers function (test.use = “MAST”, latent.vars = c(“nCount_RNA”, “nFeature_RNA”)) in Seurat v4.0. Only G1 cells were used in differential expression analysis to avoid confounding of cycling cells. Cell cycle phase was annotated with CellCycleScoring function in Seurat.

We used the fgsea^41^ v1.24.0 package to conduct gene set enrichment analysis with Hallmark gene sets (n = 50) downloaded from MSigDB^34^. We set the following parameters for the gene set enrichment analysis: nperm = 1000, minSize = 15, maxSize = 500.

#### Statistical analysis and visualization

Statistical tests and visualization were performed with R (v.4.2) package ggpubr (v.0.5.0) and ggplot2 (v.3.4).

## Data availability

Processed data containing input and output of TreeAlign have been deposited in Zenodo (https://doi.org/10.5281/zenodo.7517412).

## Code availability

The code is publicly accessible on a GitHub repository (https://github.com/AlexHelloWorld/TreeAlign), which implements TreeAlign and describes how to generate simulated datasets.

## Acknowledgements

This project was funded in part by Cycle for Survival supporting Memorial Sloan Kettering Cancer Center. SPS holds the Nicholls Biondi Chair in Computational Oncology and is a Susan G. Komen Scholar. This work was funded in part by the Cancer Research UK Grand Challenge Program to SPS [C42358/A27460], a National Institutes of Health Center for Excellence in Genome Sciences grant RM1-HG011014 and the NCI Cancer Center Core Grant P30-CA008748.

---

**Extended Data Fig. 1:**
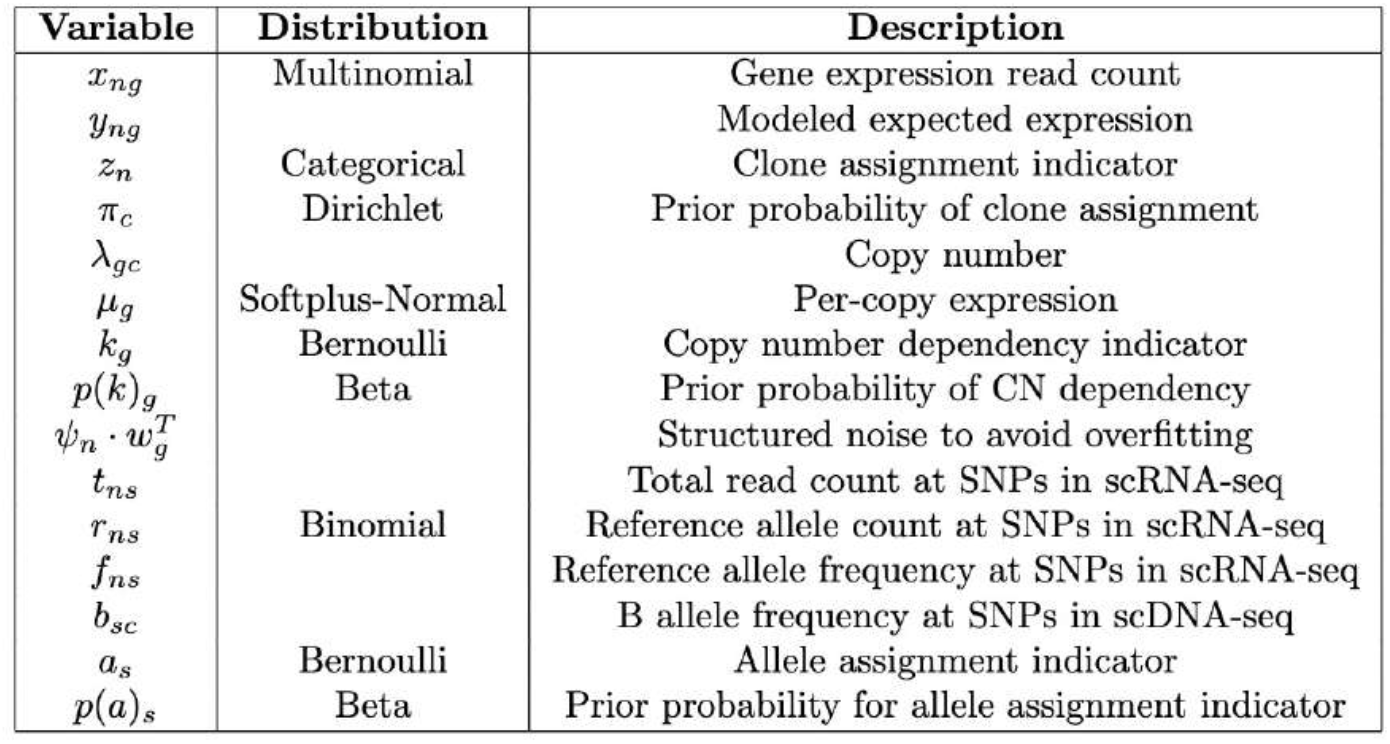
Random variables and data in TreeAlign. Descriptions and prior distributions of random variables and data in TreeAlign model.

**Extended Data Fig. 2:**
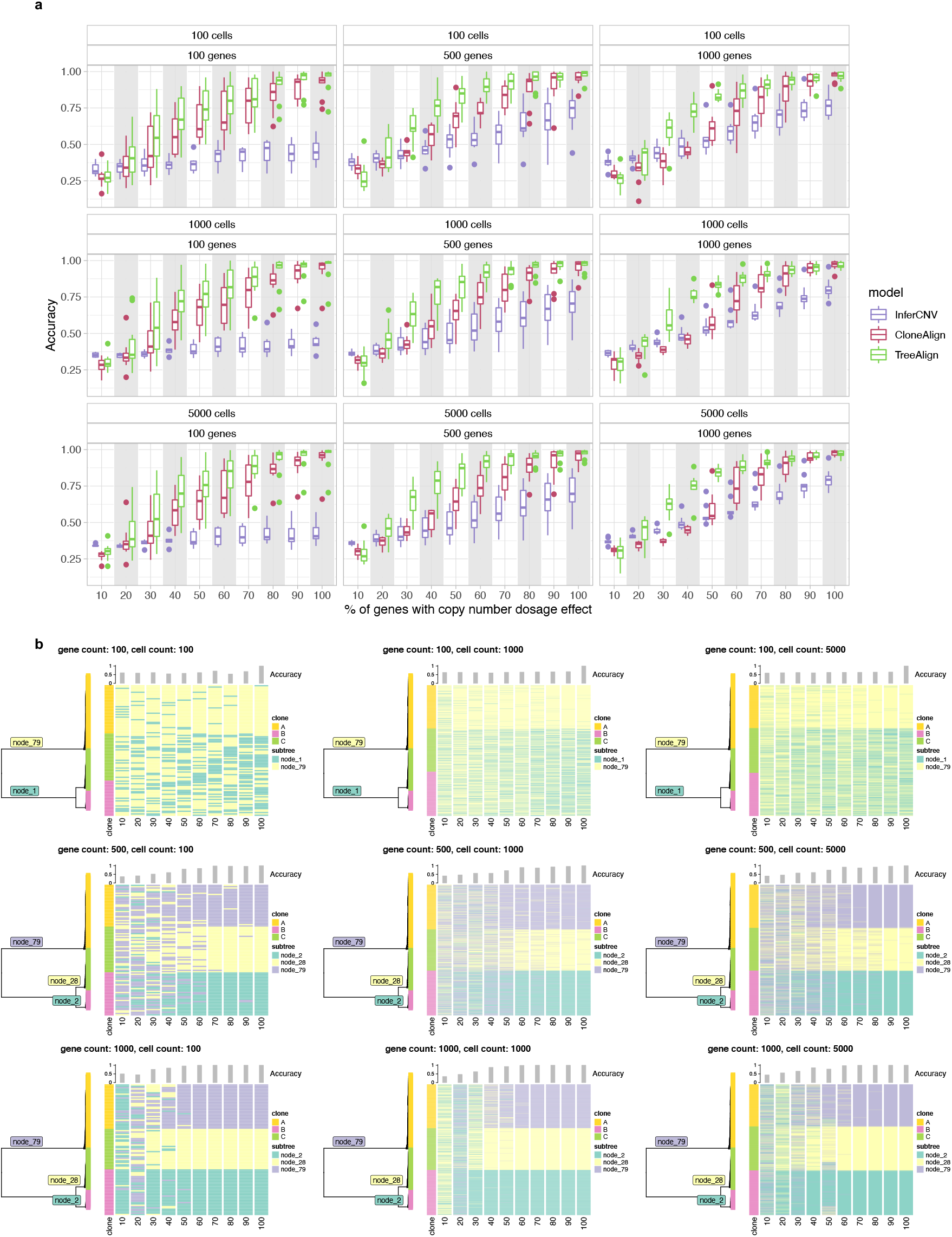
Clone assignment accuracy of TreeAlign in simulated datasets. Heat maps (right) showing clone assignment of simulated datasets by TreeAlign. **a**, Accuracy of clone assignment for TreeAlign, CloneAlign and InferCNV in simulated scRNA datasets as a function of varying proportions of genes with CN dosage effects. Panels represent datasets with different numbers of cells and genes. **b**, Phylogenetic trees (left) constructed with scDNA-data from SPECTRUM-OV-081 along with Heat maps (right) showing clone assignment of simulated datasets by TreeAlign.

**Extended Data Fig. 3:**
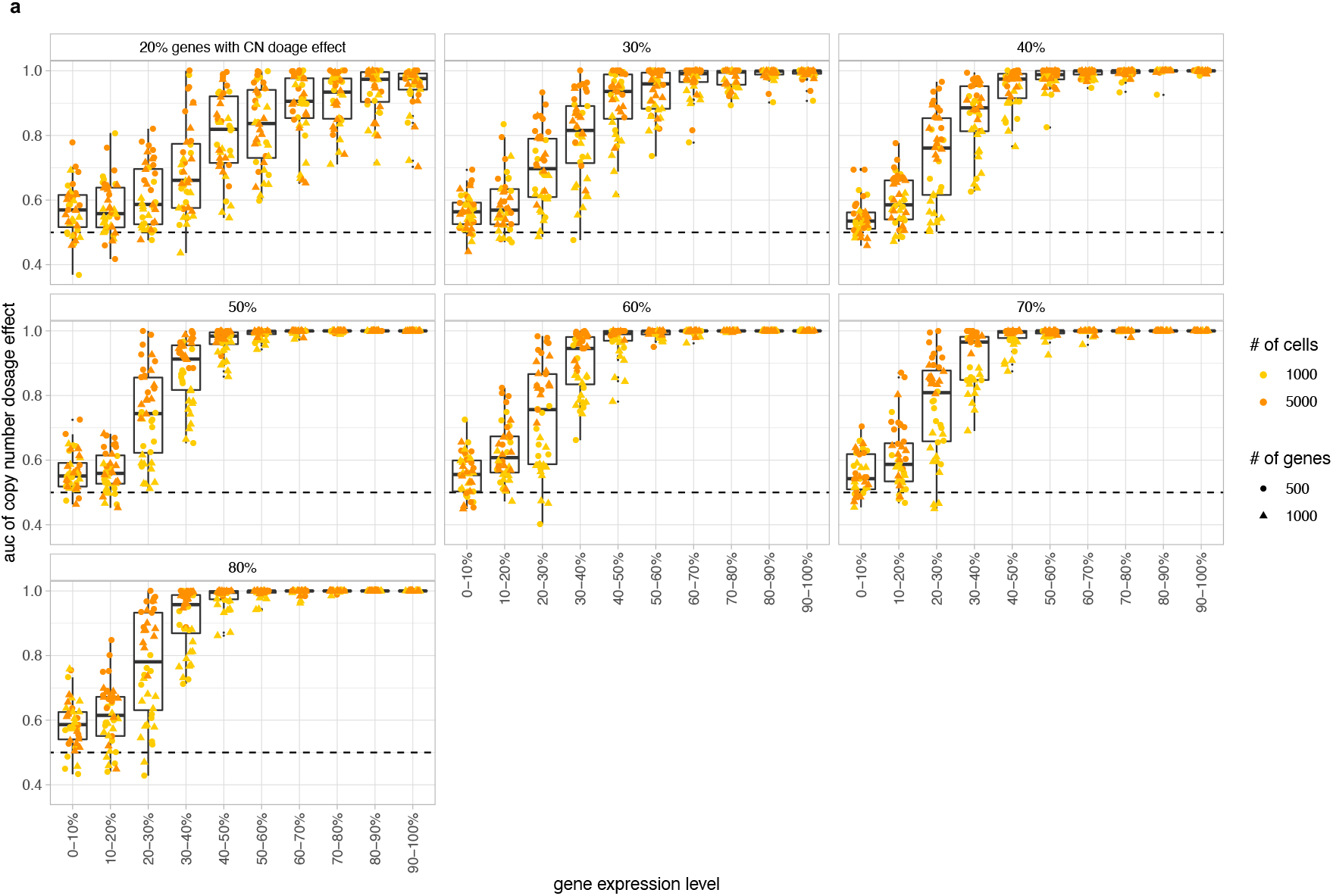
Dosage effect prediction of TreeAlign in simulated datasets. **a**, AUC of CN dosage effect *p*(*k*) predicted by TreeAlign as a function of gene expression level. Panels represent simulated datasets with varying gene dosage effect frequencies.

**Extended Data Fig. 4:**
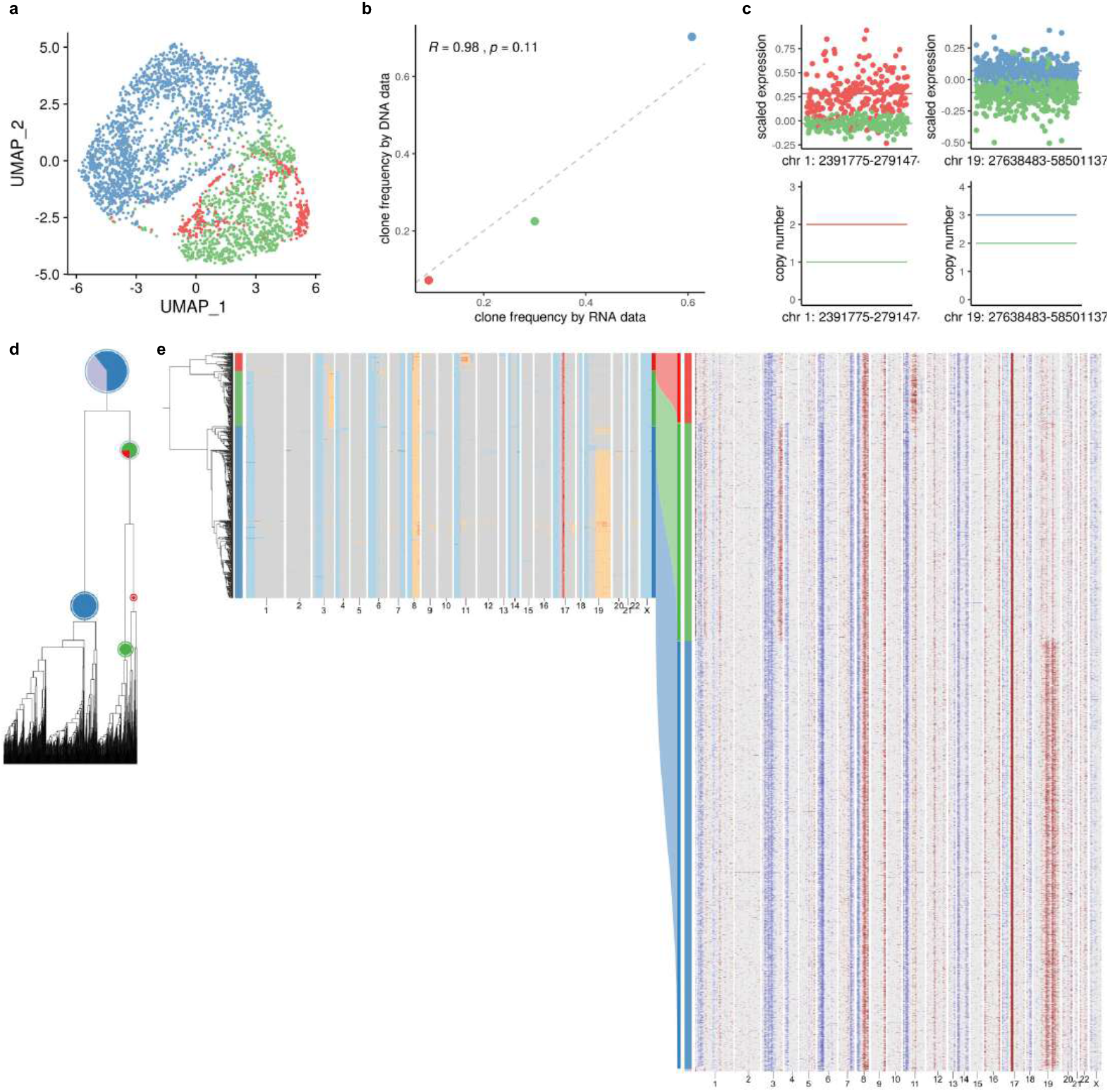
TreeAlign assigns expression profiles of NCI-N87 to phylogeny. **a**, UMAP plot of scRNA-data from gastric cell line NCI-N87 colored by clone labels assigned by total CN TreeAlign. **b**, Clone frequencies of NCI-N87 estimated by scRNA-data (x axis) and scDNA-data (y axis). **c**, Scaled expression and copy number profiles for regions on chromosome 1 and 19 as a function of genes ordered by genomic locations. **d**, Phylogenetic tree constructed with scDNA-data. **e**, Phylogenetic tree constructed with scDNA-data along with pie charts showing how TreeAlign assigns cell expression profiles to subtrees recursively. The pie charts are colored by the proportions of cell expression profiles assigned to downstream subtrees. The outer ring color of the pie charts indicates the current subtree. Heat maps of copy number profiles from scDNA (left) and InferCNV corrected expression profiles from scRNA (right). The Sankey chart in the middle shows clone assignment from expression profiles to copy number based clones by total CN TreeAlign.

**Extended Data Fig. 5:**
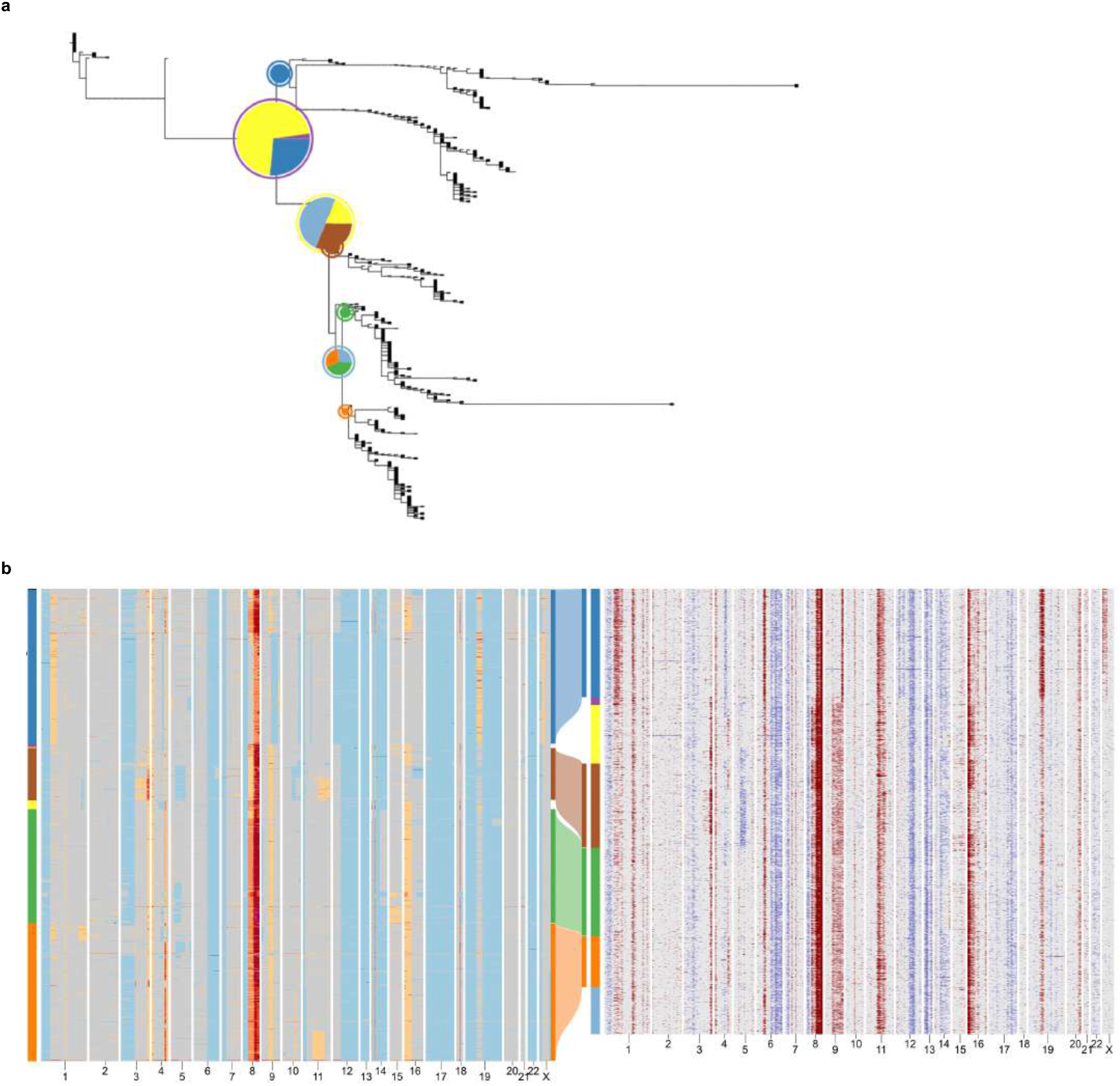
TreeAlign assigns expression profiles of patient 022 to phylogeny constructed with Sitka^38^. **a**, Phylogenetic tree constructed with scDNA-data using Sitka. Pie charts illustrate how TreeAlign assigns cell expression profiles to subtrees recursively. **b**, Heat maps of copy number profiles from scDNA (left) and InferCNV corrected expression profiles from scRNA (right). The Sankey chart in the middle shows clone assignment from expression profiles to CN-based clones characterized with Sitka.

**Extended Data Fig. 6:**
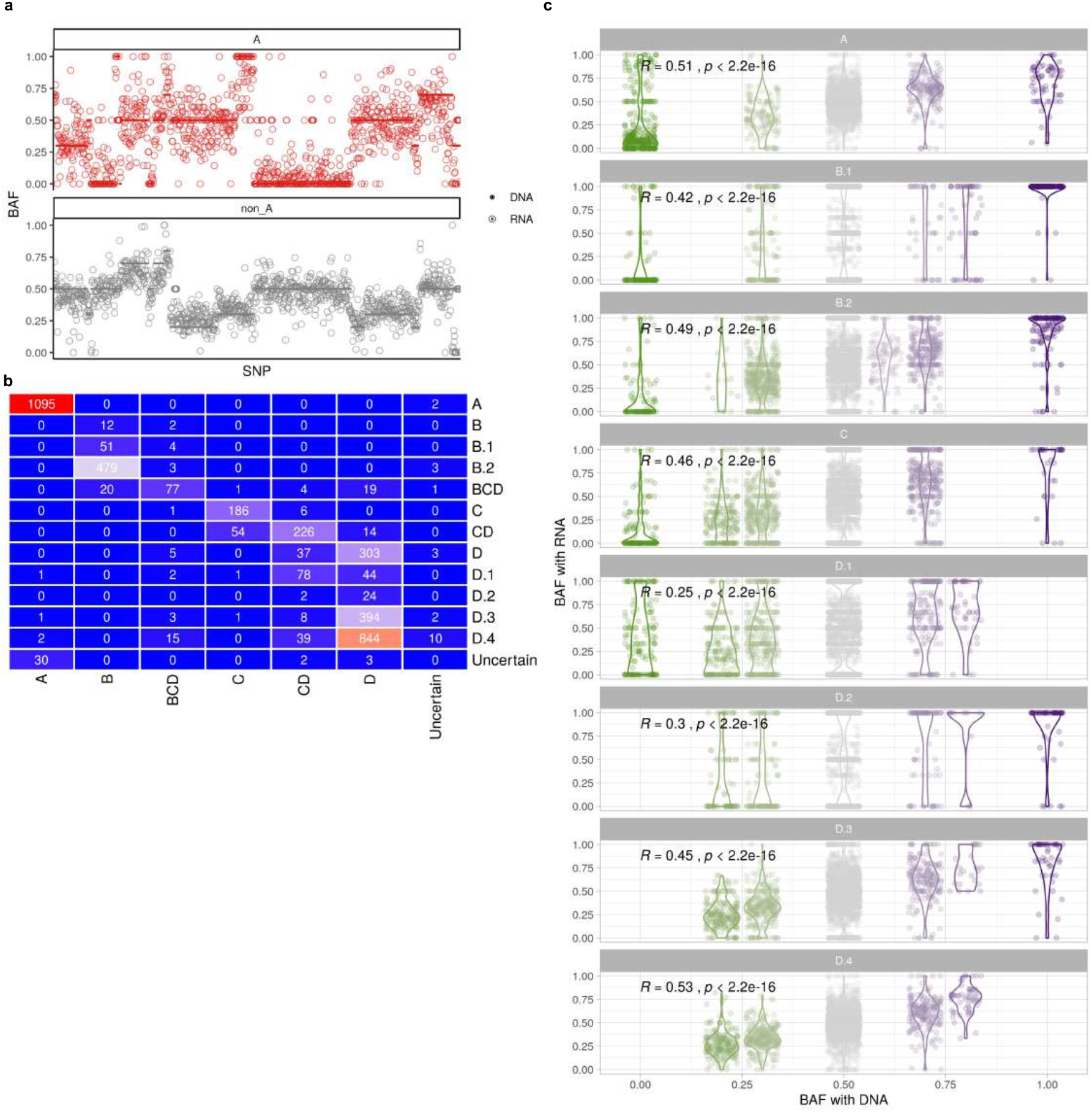
Allele-specific information contributes to clone assignment. a, BAF of heterozygous SNPs estimated from scRNA-data and scDNA-data for clone A and other clones (clone B - C) in patient 022 (ordered by gene location along chromosome). b, violin plot of BAF in SPECTRUM-OV-022 (Wilcoxon signed-rank test). **b**, Confusion matrix comparing clone assignment between total CN TreeAlign and integrated TreeAlign for patient 022. **c**, Correlation between BAF estimated with scRNA and DNA in patient 022 subclones (Wilcoxon signed-rank test).

**Extended Data Fig. 7:**
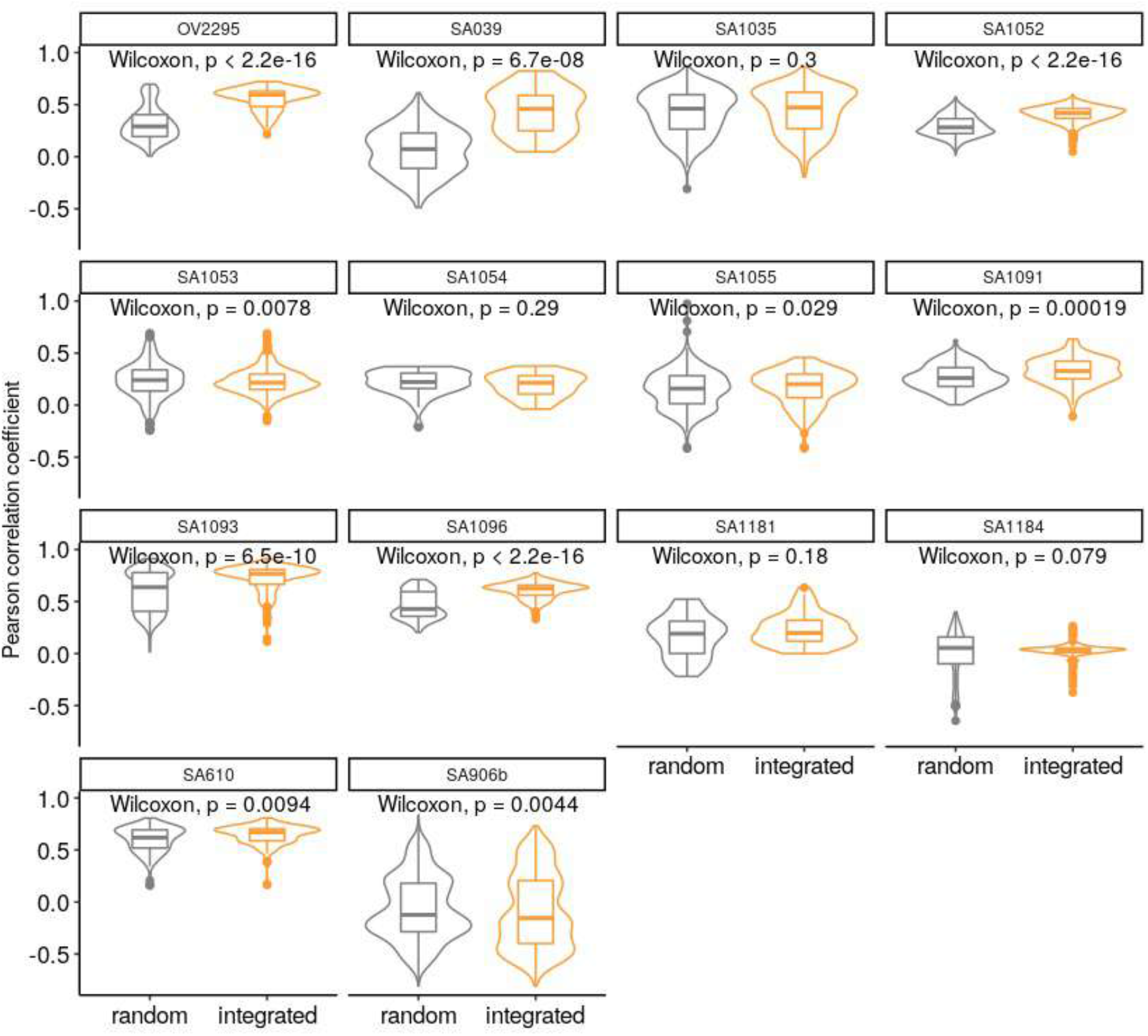
Integrated TreeAlign has improved clone assignment performance compared to total CN TreeAlign. Distribution of Pearson correlation coefficients (R) between scDNA estimated total copy number and InferCNV corrected expression for unassigned cells from total CN model. Left, correlation distribution calculated by comparing InferCNV profiles to CN profiles of a random subclone; Right, correlation distribution calculated by comparing InferCNV profiles to CN profiles of subclones assigned by integrated TreeAlign. Each panel represents results from a tumor sample/cell line.

**Extended Data Fig. 8:**
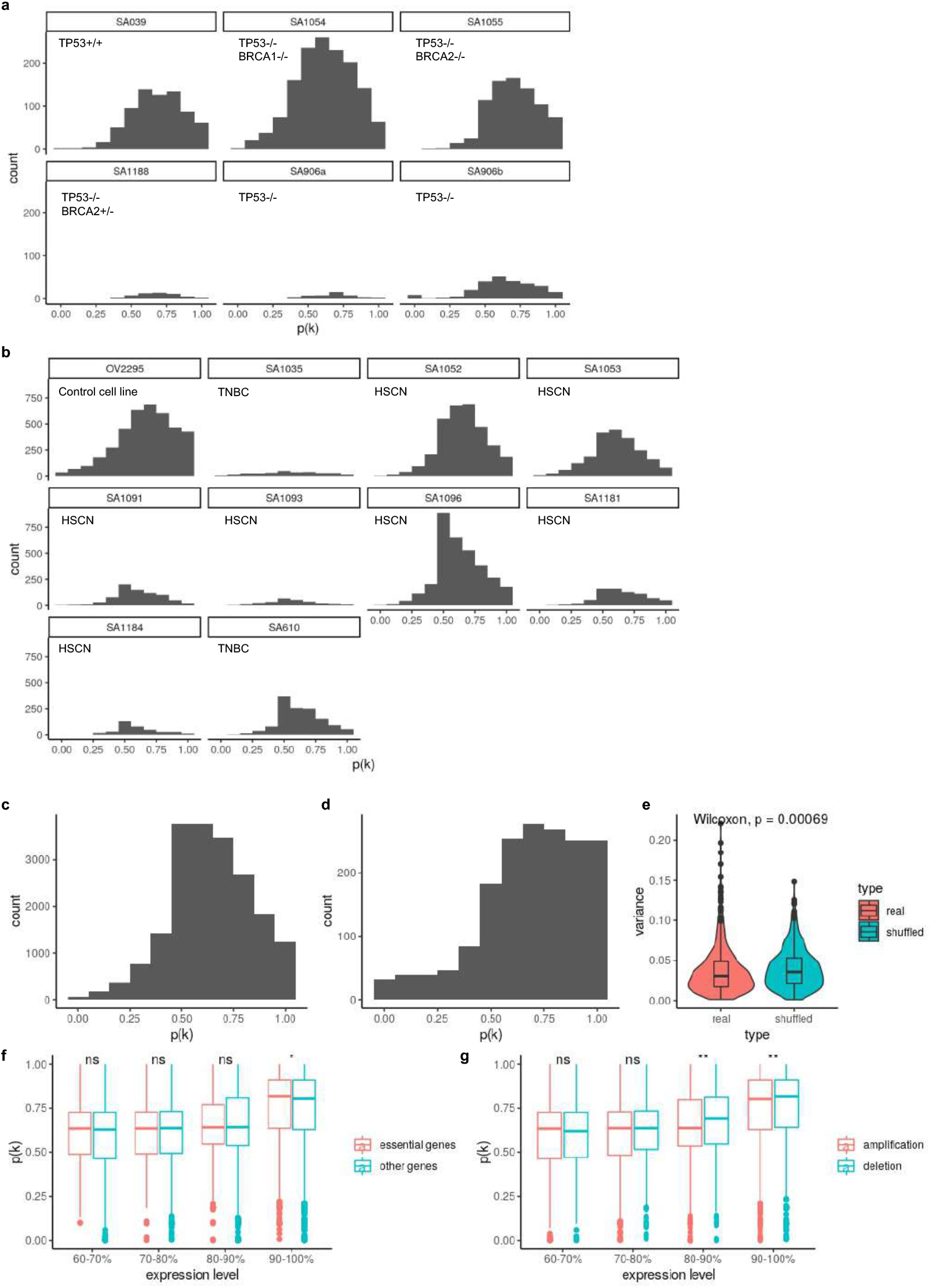
Distribution of *p*(*k*) in tumors and cell lines. **a**, Distribution of ***p***(***k***) hTERT-184 cell lines. **b**, Distribution of ***p***(***k***) in ovarian cancer control cell line OV2295, primary HGSC and TNBC tumors. **c**, Distribution of ***p***(***k***) across primary tumors and cell lines in and (b). **d**, Distribution of ***p***(***k***) in patient 022. **e**, Variance of ***p***(***k***) of the same gene between patients compared to variance of randomly shuffled ***p***(***k***). **f, *p***(***k***) distribution as a function of gene essentiality^43^ in gene groups with different expression levels. Only high expression genes (top 40%) are shown. **g, *p***(***k***) distribution between genes located in amplifications and deletions. Only high expression genes (top 40%) are shown.

**Extended Data Fig. 9:**
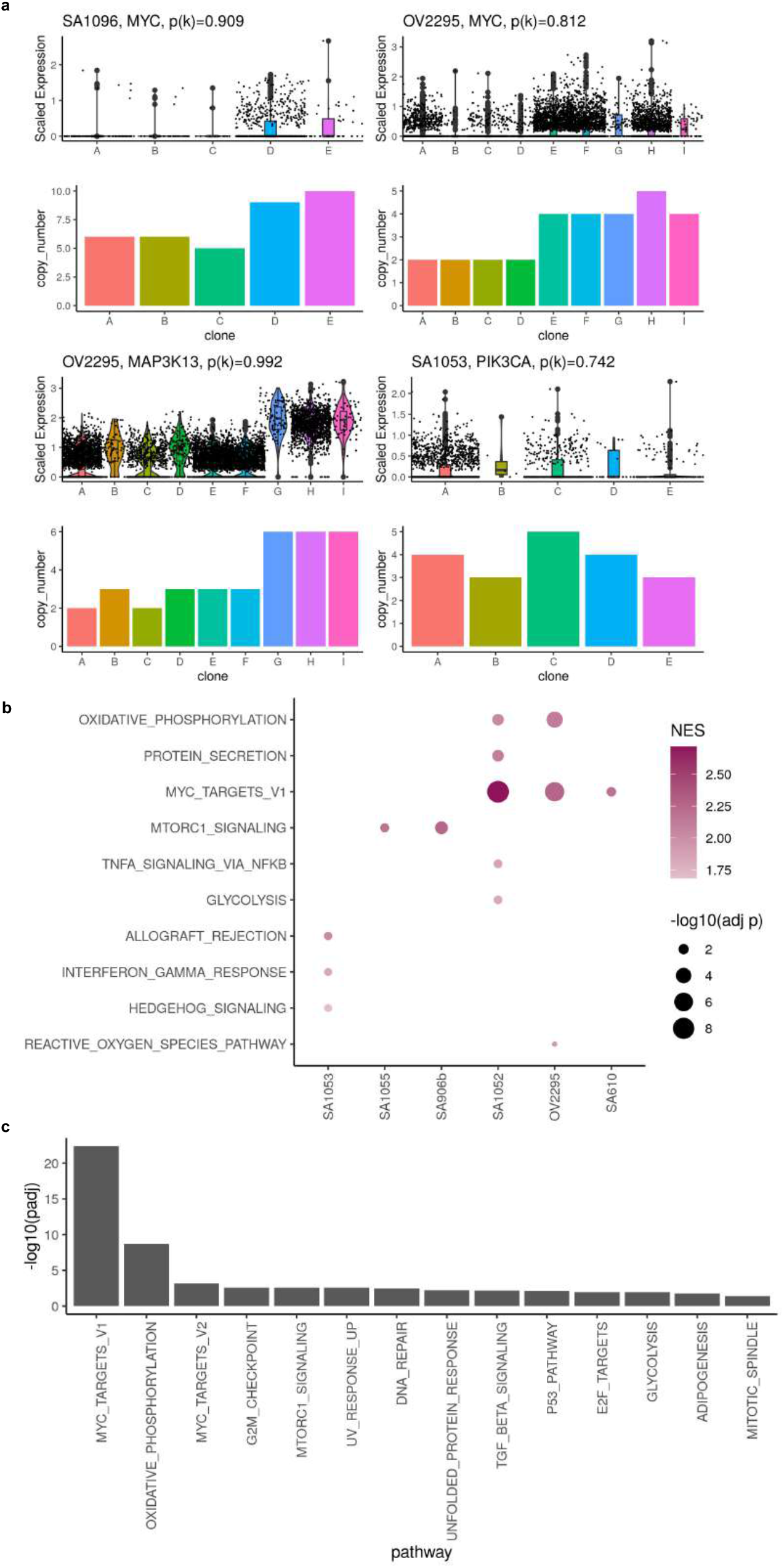
Gene set enrichment analysis of low *p*(*k*) genes. **c**, Example of genes with high level amplifications and high CN dosage effects. **b**, Dot plot showing significantly enriched pathways in low ***p***(***k***) genes. **b**, Significantly enriched pathways in low ***p***(***k***) genes from all primary tumors and cell lines. ***p***(***k***) from all samples were combined before performing gene set enrichment analysis.

**Extended Data Fig. 10:**
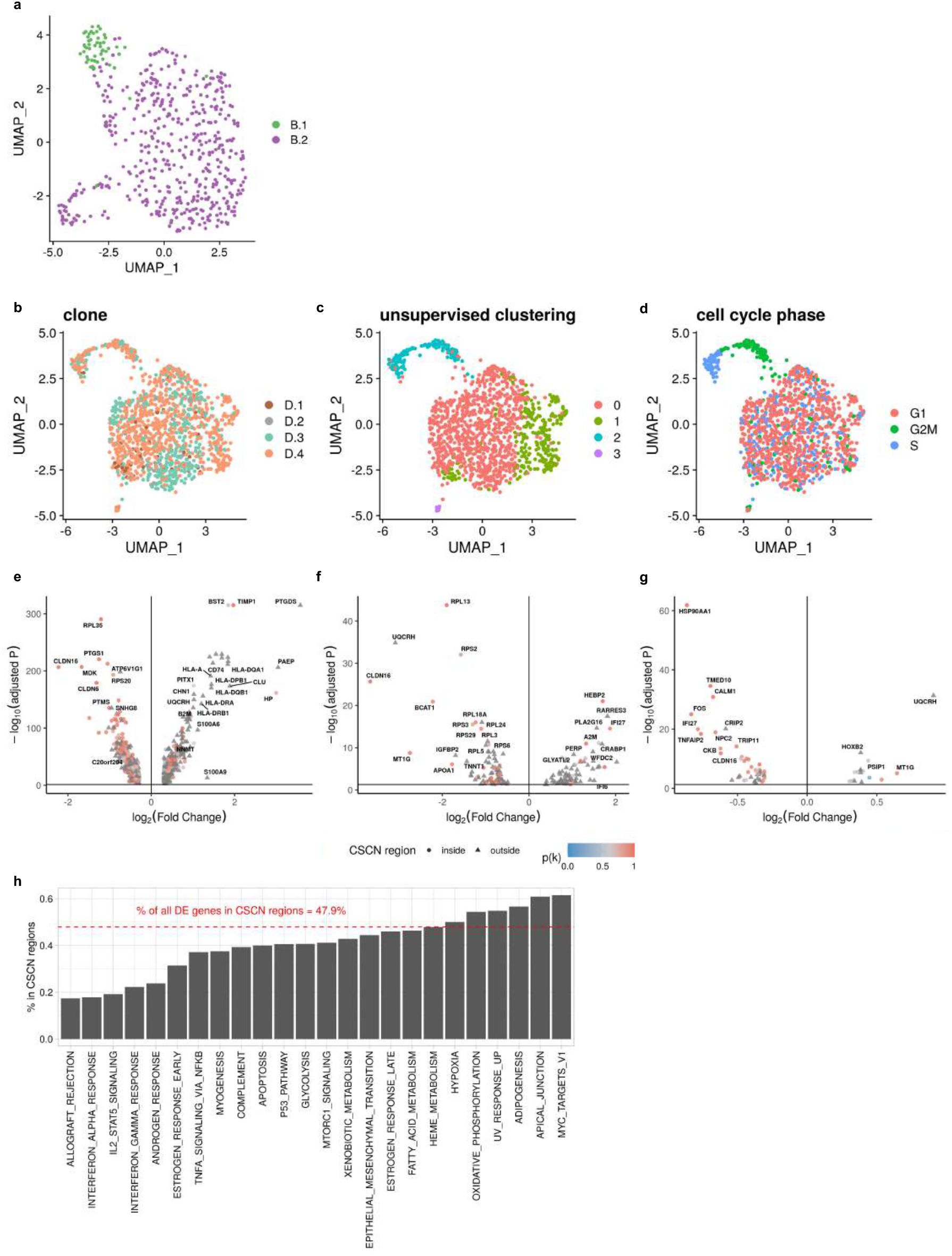
Differentially expressed genes between subclones in patient 022. **a**, UMAP plot of expression profiles of clone B.1 and B.2 in patient 022. **b**, UMAP plot of expression profiles of clone D.1, D.2, D.3 and D.4 in patient 022 colored by clone assignments. **c**, UMAP plot of expression profiles of clone D in patient 022 colored by Louvain unsupervised clustering. **d**, UMAP plot of expression profiles of clone D in patient 022 colored by cell cycle phase. **e**, Differentially expressed genes between clone A and clone B - D. **f**, Differentially expressed genes between cells in clone B.1 and B.2. **g**, Differentially expressed genes between cells in clone D.4 and D.1 - D.3. **h**, Frequencies of DE genes in CSCN regions summarized by Hallmark pathways.

**Extended Data Fig. 11:**
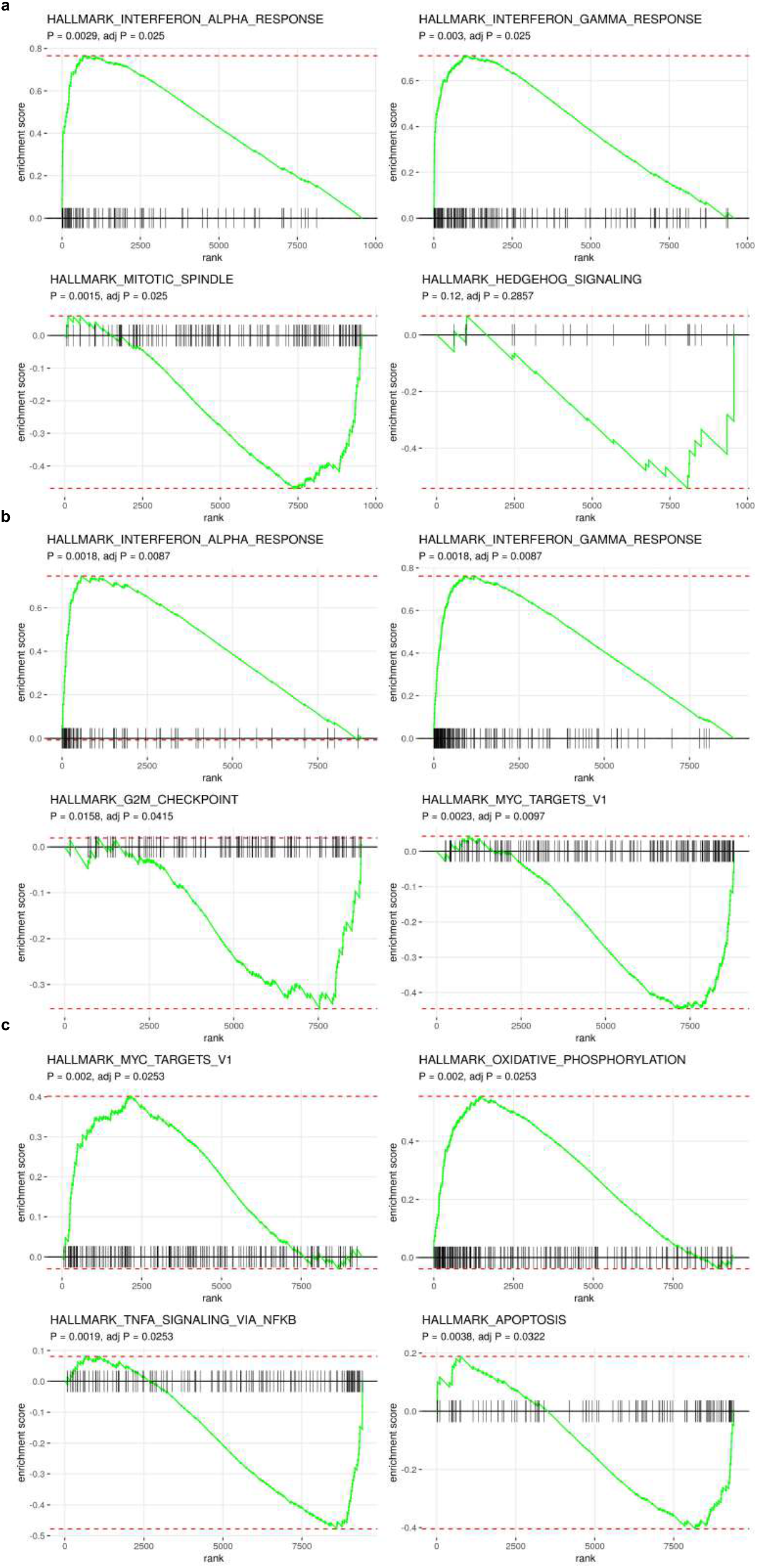
Examples of enriched and depleted pathways in patient 022 subclones. **a**, Enriched and depleted pathways in clone A compared to other clones in patient 022. **b**, Enriched and depleted pathways in clone B.1 compared to clone B.2. **c**, Enriched and depleted pathways in clone D.4 compared to the rest of cells in clone D.

